# Neural Correlates of Trial Outcome Monitoring during Long-term Learning in Primate Posterior Parietal Cortex

**DOI:** 10.1101/2025.08.05.668637

**Authors:** Ziang Liu, Zhuangyi Jiang, Li Shi, Fang Fang, Shiming Tang, Yang Zhou

## Abstract

Monitoring behavioral outcomes is crucial for optimizing behavior and learning new tasks. However, how the brain monitors outcome information during long-term learning remains largely unknown. Using two-photon calcium imaging in behaving monkeys, we tracked neuronal activity in area 7a of the posterior parietal cortex (PPC), an important associative cortex that has not been typically implicated in associative learning (AL), while the macaque monkeys learned new visual-motor associations over multiple days. We found robust neuronal representation of salient behavioral outcomes (correct vs. incorrect) which closely correlated with the monkeys’ learning behavior. Furthermore, outcome representation significantly reorganized when monkeys transitioned to learn novel associations, followed by gradual evolution over subsequent learning days, a process constrained by the functional connectivity among neurons within local network. These suggest a substantive role for primate PPC in long-term AL through monitoring trial outcome, and indicate a principle for long-term learning: network connectivity constrains the evolution of neural encodings.

## Introduction

The ability to learn appropriate motor responses to different environmental stimuli is essential for an animal’s survival in complex environment. During AL, the effectiveness of establishing new sensorimotor associations relies on the brain’s ability to monitor the positive and negative outcomes of our action choices during individual learning trials^1-3^. This outcome monitoring allows brain to evaluate the action consequences, reinforce relevant associations, and guide adaptive changes. Theoretically, brain regions central to outcome monitoring during AL should fulfill several criteria. First, they should exhibit significant encoding of trial outcomes (correct vs. incorrect) following the completion of each trial, with outcome signals that go beyond mere reward reception to reflect higher-level cognitive processing. Second, the strength or pattern of outcome encoding should correlate with the animal’s learning performance or strategy and adapt dynamically to the learning context. Third, such outcome representations should contribute causally or functionally to animal’s subsequent behavior during the learning process.

The prefrontal cortex, cingulate cortex, and other subcortical areas have been implicated in representing trial outcomes during sensorimotor AL^3-10^, as their neural activity reflects the correctness of animals’ action choice during short-term AL. These outcome representations were not merely reflective of reward reception; some were correlated with the animals’ behavior performance during leaning^6^. However, how the brain monitors behavioral outcomes remains elusive, as the previous studies only partially met one or two criteria. Furthermore, acquiring complex sensorimotor skills often requires gradual learning over periods spanning days or longer, but previous primate studies have primarily focused on the changes of neuronal activity during relative short period of minutes to hours during the learning process^6,11-17^. Since no study has tracked the evolution of neuronal encoding of the same population of neurons in the primate brain over different learning days, our understanding of the mechanisms underlying long-term AL is highly limited. Key questions regarding outcome monitoring remain untested, particularly during long-term AL: How does the neuronal representation of trial outcomes correlate with the animals’ behavior to potentially influence learning? How does the outcome representation change throughout the learning process and what factors govern these changes?

Previous studies have highlighted the fronto-striatal circuit as the central hub for sensorimotor AL^15,17-24^, with less emphasis placed on the roles of other brain areas. However, changes in neuronal activity were observed in other brain areas both during and after sensorimotor AL^12,25-28^, suggesting the engagement of a potentially broader brain network. The PPC, a crucial association cortex positioned at the intermediate stage of sensorimotor transformation, has been implicated in various cognitive functions^29-34^, but was usually not implicated in AL. The PPC has been shown to play important roles in representing stimulus-stimulus association and sensorimotor association after long-term learning^26,35,36^, implying potential involvement in sensorimotor AL. Specifically, neuronal activity in the lateral parietal cortex has been reported to reflect outcome information (rewarded vs. non-rewarded) during a decision task^37^. Nonetheless, the exact role of PPC in long-term AL remains largely unexplored.

Area 7a, a subregion of the PPC, occupies the highest level of the visual processing hierarchy in the parietal lobe^38^. Traditionally, it has been implicated in diverse cognitive functions, including spatial attention, multisensory integration, motor planning, coordinate transformations, and other higher-order visuospatial processes^33,34,39-44^. However, its core computational roles remain debated. Notably, Area 7a exhibited direct connections with regions involved in learning and memory, such as prefrontal cortex and medial temporal lobe^38,45,46^, as well as with structures engaged in sensorimotor transformation, such as oculomotor areas^47^, suggesting a potential role in sensorimotor AL. Despite this, direct empirical evidence for its involvement in AL—and the mechanisms through which it may contribute—remains unexplored.

In this study, we investigated the role of primate PPC in long-term AL by using two-photon calcium imaging to record the activity of a large population of area 7a neurons, while monkeys were learning image-saccade associations. We found that about half of these 7a neurons distinctly represented the trial outcome and its recent history, and this outcome representation closely correlated with the monkeys’ learning behaviors. Interestingly, outcome representation within 7a was primarily sensitive to outcome switches, and potentially drive the exploration behavior in the subsequent trial. Remarkably, by tracking these neurons over the course of long-term learning, we found that the neuronal representation of trial outcome substantially reorganized when monkeys transitioned from exploiting familiar association to exploring novel association, and then gradually evolved in a manner contingent on the learning content during the acquisition of new associations learned over multiple days. Critically, this evolution of outcome representation was constrained by the local network structure, as reflected by functional connectivity between neurons. Together, our findings provide new insights into the neural mechanisms of outcome monitoring during long-term learning, highlighting a substantive role for primate PPC in AL, a function traditionally less appreciated.

## Results

### Representation of trial outcomes in area 7a

We trained two Macaque monkeys to perform an image-saccade association learning (ISAL) task (**Figure 1a**), in which monkeys had to choose one of two saccade targets based on a presented visual image. Monkeys learned to associate different visual images with corresponding saccade directions based on the trial outcome – receiving juice reward or a brief time-out punishment after correct or incorrect saccade choices, respectively. The reward was delivered immediately after disappearance of the saccade target in correct trials, ensuring that the trial outcome was swiftly apparent in both correct and incorrect trials. Each time the monkeys mastered two image-saccade associations (accuracy 0.89±0.05, mean±STD), they were given two novel associations in subsequent days (**Figure 1b**), with the familiar associations no longer being tested at this stage of learning. While the monkeys learned new image-saccade associations, the saccade targets themselves remained constant. The monkeys’ performance accuracy substantially decreased on the first day when a new association was introduced (0.61± 0.12, **Figure 1c-d**), and then gradually rose to a high level within a few days (3.9 ± 1.3 days). We employed a dynamic model, PsyTrack^48^, to fit the monkeys’ performance accuracy across learning each pair of associations (**Figure 1c, e-f**). This model describes choice behavior at the resolution of single trials, allowing us to quantify the evolution of animal’s decision strategies (represented by different weights) across the learning course. It revealed a progressive increase in the weight attributed to visual stimuli, coupled with a gradual decrease in choice bias when learning new associations, consistent with the monkeys gradually learned the new associations.

**Figure 1.**
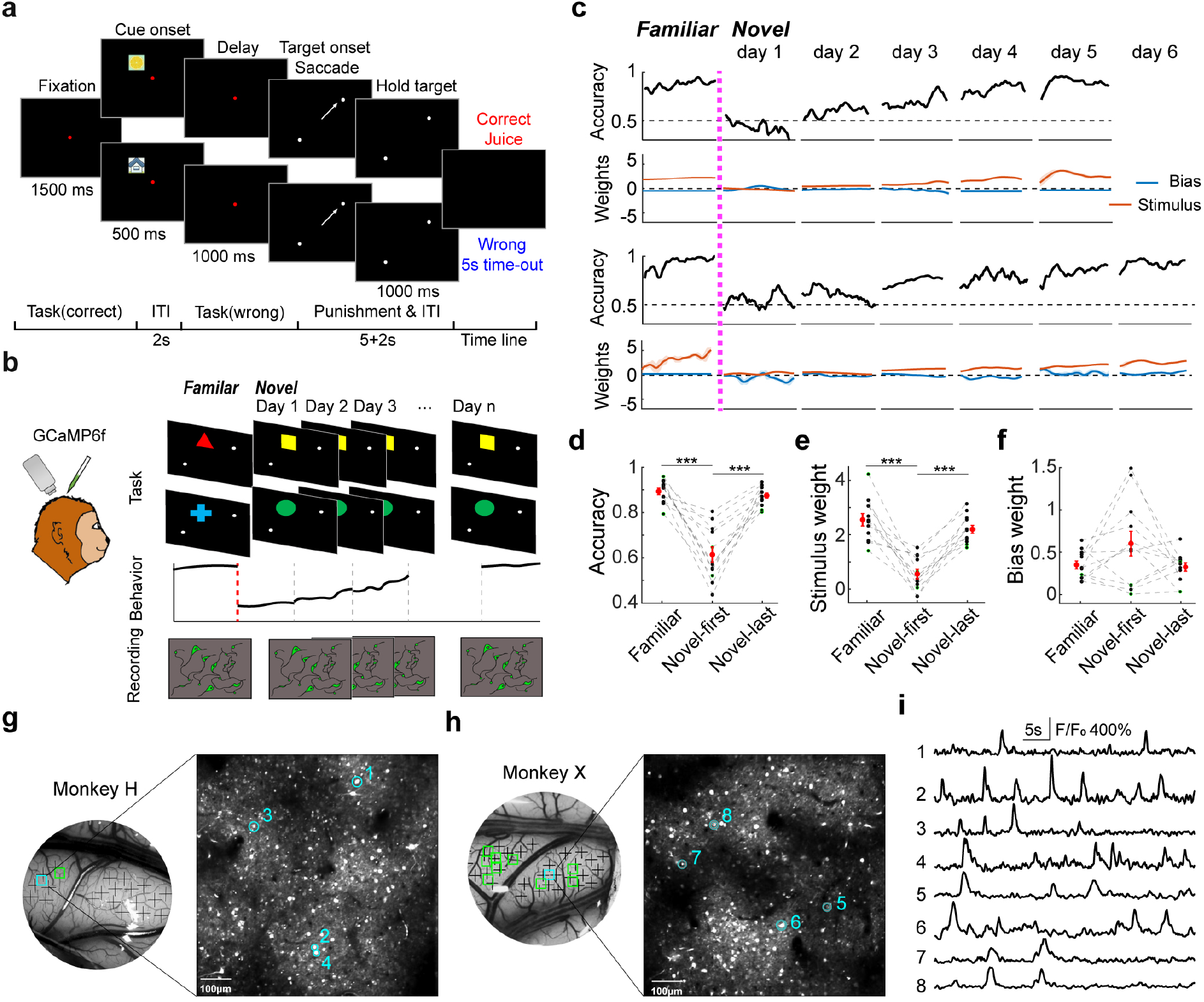
Two-photon calcium imaging of macaque PPC during sensorimotor AL. **a**. Schematic illustration of the image-saccade association learning (ISAL) task. **b**. Experiment design. Neural activities of individual 7a neurons were recorded using two-photon calcium imaging while monkeys learned image-saccade associations. Across each learning course, we monitored the same neurons, beginning from the day when monkeys were tested with familiar associations until they successfully acquired a pair of novel associations. **c**. Monkeys’ behavior performance during two example learning courses. The pink vertical dashed line marks the transition when monkeys began to learn novel associations. The first and second rows depict the evolution of the monkey’s performance accuracy over different days in one learning course, accompanied by the corresponding stimulus weight and bias weight derived from fitting the ‘PsyTrack’ model. The third and fourth row display the data from the other example learning course. Shaded region denotes the 95% confidence interval. **d**. Comparisons of each monkey’s averaged performance accuracy across different days in learning courses. Each point denotes one data session, with dashed lines connecting data from the same learning course. Data from two monkeys are shown in green and black separately. The red dot signifies the average results over 12 learning courses. Error bars represent SEMs. **e-f**. Comparisons of the stimulus weight and bias weight. **g-h**. Two example FOVs, each from one monkey respectively. The brain areas under the recording chambers are marked with the sites where the virus was injected (black crosses). The cyan rectangles denote the locations of the example FOVs. The small rectangles denote the FOVs. **i**. The normalized fluorescent intensity traces (F/F_0_) of 8 example neurons from the two FOVs. (Novel, first: Novel cue, first learning day. Novel, last.: Novel cue, the last learning day; ***: p<0.001, paired t-test).

To monitor the neuronal activity during sensorimotor AL, we injected AAV to induce GCaMP6f expression in neurons at a depth of ∼400 μm within PPC area 7a, allowing us to simultaneously record the activity of hundreds of neurons over multiple days (**Figure 1b, g-h**). The recordings spanned depths of 200–300 μm beneath the arachnoid membrane, primarily corresponding to layers 2–3. We recorded activity from a total of 27,511 single neurons with a good signal-to-noise ratio from two monkeys (see Methods). We found that these 7a neurons encoded trial outcome after the saccade target disappeared (ranksum test, P < 0.01, Bonferroni corrected, see Methods): some neurons exhibited stronger responses following incorrect saccade choices (error neurons, ENs; **Figure 2a**); others did the opposite (correct neurons, CNs: **Figure 2b**). To quantify the strength of neuronal encoding related to trial outcomes, we used one-way ANOVA to calculate the unbiased fraction of explained variance (FEV) in the neuron’s activity attributable to the trial outcome (see Methods). **Figure 2c-d** show an example of field-of-view (FOV) within which most neurons (92%) showed significant outcome selectivity. Furthermore, ENs outnumbered the CNs (92 vs. 51), and there was no obvious spatial clustering for either type of outcome-selective neuron.

**Figure 2.**
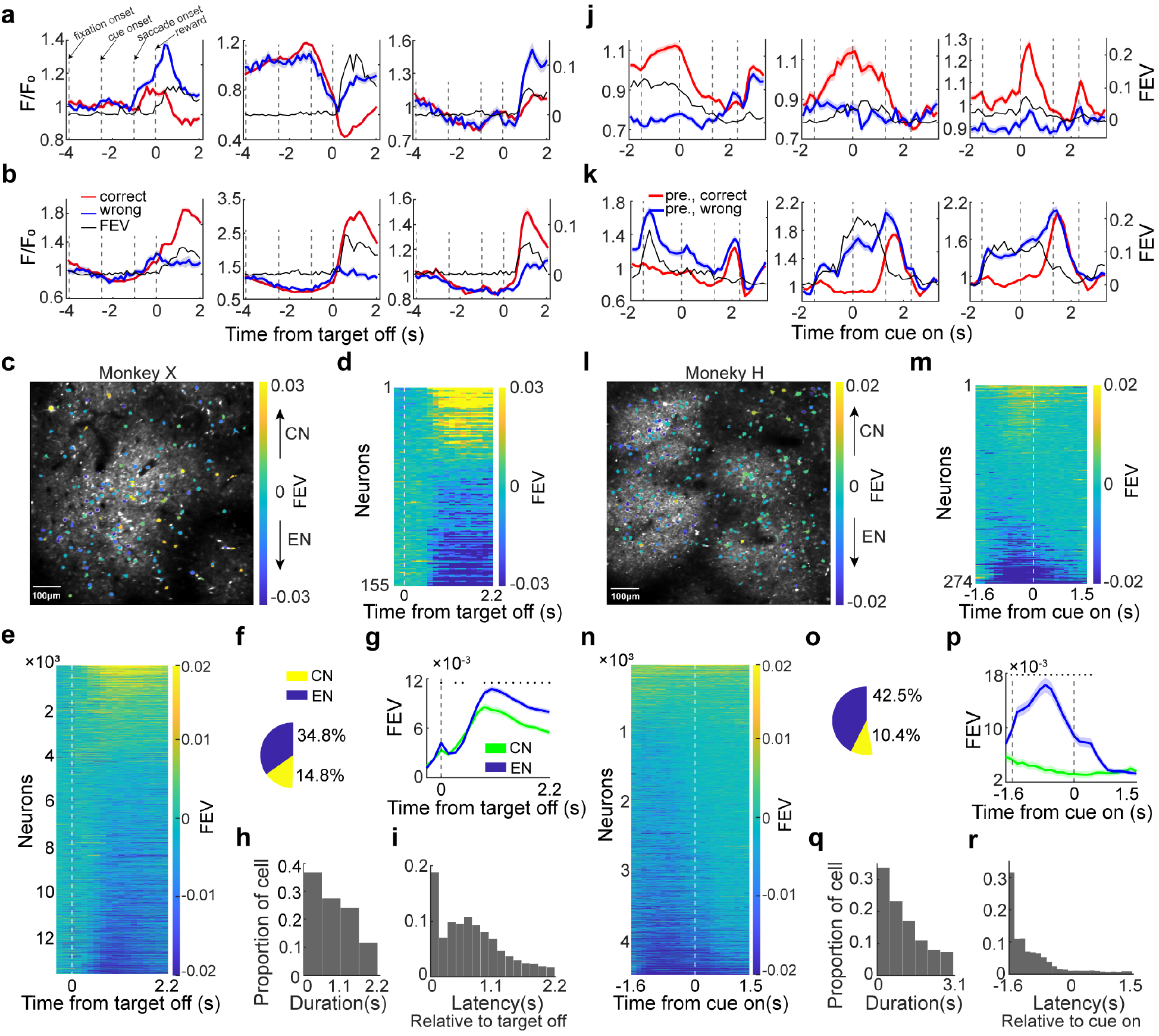
7a neural activity represents trial outcome during AL. **a-b**. Example outcome-selective neurons. The normalized activity (F/F_0_) and the magnitude of outcome selectivity (black, quantified by FEV analysis) of 3 error-preferred neurons (**a**, EN) and 3 correct-preferred neurons (**b**, CN) are displayed over time relative to saccade target offset (0 s). Other vertical dashed lines (from left to right) indicate the approximate timing of fixation onset, cue onset and saccade onset. **c**. An example FOV with identified ROIs. The color bar marks the averaged strength (FEV value) and preference of outcome selectivity. **d**. Outcome-selectivity for all identified neurons in the example FOV. **e**. Outcome selectivity for all identified outcome-selective neurons (13,594 of 27,511) across all FOVs. **f**. Percentages of CNs and ENs among all identified neurons. **g**. The averaged magnitude of outcome selectivity were shown separately for both CNs and ENs. Shaded regions denote SEMs. Stars denote the time points with significant difference between CNs and ENs (P < 0.01, t test). **h**. Duration of outcome encoding for all outcome-selective neurons. Each time window spans 0.54s. 63% neurons exhibited significant outcome encoding in two or more time windows. **i**. Latency of outcome selectivity relative to target offset for all the outcome-selective neurons. **j-r**. 7a neurons represented the trial outcome in the previous trial (outcome history). **j-k**. Example neurons. Activities were sorted according to the trial outcome in the previous trial. **l-m**. An example FOV, shown in the same format as (c-d). **n-r**. The PO encodings of all selective neurons (4489 of 8503 neurons from monkey H).

Population results from both monkeys were well represented by the phenomenon observed in the example FOV (**Figure 2e-i**, **Supplementary Fig. 1-2**). About half of the identified neurons exhibited significant encoding of trial outcome, with greater prevalence to negative outcomes (14.8% CNs, 34.8% ENs, **Figure 2e-f**). Even though both CNs and ENs showed persistent outcome encoding on the population level, ENs exhibited significantly greater outcome selectivity than CNs (**Figure 2g**). Most outcome-selective neurons (63%) exhibited sustained outcome encoding beyond 1 second (**Figure 2h)**. We did not detect spatial clustering for the outcome encoding in the majority of the FOVs (**Supplementary Fig. 3**). Moreover, although the outcome encoding in 7a spans a wide-ranging latency throughout the inter-trial interval (**Figure 2i**), it cannot be attributed to differences in monkeys’ eye positions or eye movement-related behaviors between correct and error trials during this period. First, there is no significant correlation between the strength of outcome encoding and the monkeys’ eye positions during the inter-trial interval (**Supplementary Fig. 4a-b**). Second, both monkeys exhibited similar saccade behaviors between correct and error trials during the inter-trial interval for the majority of recording sessions, including direction (82.91%), frequency (62.39%), and amplitude (56.41%), and the difference of saccade behavior between correct and error trials did not significantly contribute to 7a neuronal activity (**Supplementary Fig. 4c**). Third, although the monkeys’ blink frequencies during the inter-trial interval differed between correct and error trials in more than half of the recording sessions (64.1%), 7a neural activity did not significantly correlated with the frequency of eye blinks during the inter-trial interval (**Supplementary Fig. 4c**). These results suggested a stable and sustained outcome encoding in the primate PPC, predominantly representing the negative outcomes.

We next tested whether 7a neuronal activity reflected the outcome history of the previous trials. For this analysis, we focused solely on data from one monkey due to the high frequency of fixation breaks (fix-break ratio = 0.57±0.08) during performing the task exhibited by the other monkey. This resulted in an excessive number of trials with no choice interspersed between completed trials. Using a linear regression model, we found that 7a neurons most strongly encoded the outcome of the immediately preceding trial, rather than those coming earlier (**Supplementary Fig. 5**). We therefore focused on analyzing the encoding of outcome of the preceding trial. As illustrated by the examples shown in **Figure 2j-k**, 7a neurons could show significant encoding of outcome-history during the fixation, sample and delay period of the current trial. In an example FOV (**Figure 2l-m**), and 57.8% of neurons exhibited significant encoding of the outcome history. Among these, ENs significantly outnumbered CNs (p << 0.001, Chi-square test) and neither neuron type showed significant spatial clustering. Population results were similar (**Figure 2n-r** and **Supplementary Fig. 6**), with 52.9% the identified neurons showing significant encoding of the outcome history (**Figure 2n-o**, 10.4% CNs, 42.5% ENs). The EN signal significantly surpassed that of CNs (P = 6.54e-30, Wilcoxon test) and peaked during the fixation period (**Figure 2p**). The outcome-history encoding of the majority 7a neurons predominantly emerged and localized within the fixation period of the ISAL task (**Figure 2q-r**). Data from the other monkey also exhibited significant encoding of the outcome history (**Supplementary Fig. 7**). Notably, the outcome-history encoding was not merely hysteresis (**Supplementary Fig. 8**): 39% of neurons exhibited inconsistent encoding of the trial outcome and outcome history, either showing significant encoding of only one of them or displaying significantly opposite preferences (7.3%).

We also quantified the activity of 7a neurons during other task epochs in the ISAL task (**Supplementary Fig. 9**). Consistent with previous findings, a substantial proportion of neurons encoded saccade direction, particularly during the post-saccade epoch (delay period: 20%, peri-saccadic period: 31%, post-saccadic period: 64%). The average strengths of saccade direction encoding and outcome encoding were positively correlated both within individual learning days and across learning days (**Supplementary Fig. 10a-b**). In contrast, very few neurons responded robustly to the visual cue (0.2%) or exhibited significant encoding on the image identity during the cue presentation period (<4%). Notably, the saccade direction selectivity during the delay period—which may reflect image-saccade associations—was notably modest (**Supplementary Fig. 10c**). Furthermore, pre-saccadic encoding in area 7a did not exhibit systematic changes across learning days that aligned with the monkeys’ learning performance (**Supplementary Fig. 10d**). These results suggest that area 7a is unlikely to be a central region for encoding and acquiring sensorimotor associations.

To further quantify the strength of neural encoding of current outcome, outcome history and saccade direction, we applied de-mixed principal component analysis to decompose 7a population activity into individual components corresponding to neural representations of different task variables (see Methods). This revealed significant encoding of all variables at the population level (**Supplementary Fig. 11**). Trial outcome and outcome history together explained significantly more variance of the population activity than the saccade direction did (P = 3.53e-16, t(32) = 15.18, paired t-test; history: 0.173 ± 0.022, outcome: 0.114 ± 0.010, saccade: 0.108 ± 0.011).

### Relevance for outcome monitoring

Upon detecting prominent outcome encoding in area 7a, we proceeded to assess whether it satisfies the three criteria for outcome monitoring.

Initially, we tested whether outcome signal in 7a just reflected reward reception. To test this, we first compared outcome encoding for the same neurons between in the ISAL task and a separate visual saccade task (**Supplementary Fig. 12**). In this task, monkeys performed saccades like those in the ISAL task but without learning associations, with rewards doubled or cancelled in a quarter of randomly chosen trials (see Methods). If outcome selectivity primarily correlated with reward reception, we expected similar levels of outcome selectivity between the tasks. Instead, both the proportion of significantly outcome-selective neurons and the population-level strength of the outcome encoding dramatically diminished during the visual saccade task (**Figure 3a-d**, P = 2.07e-11, F = 46.92, repeated ANOVA), suggesting that the outcome encoding in 7a during ISAL task correlated with sensorimotor AL.

**Figure 3.**
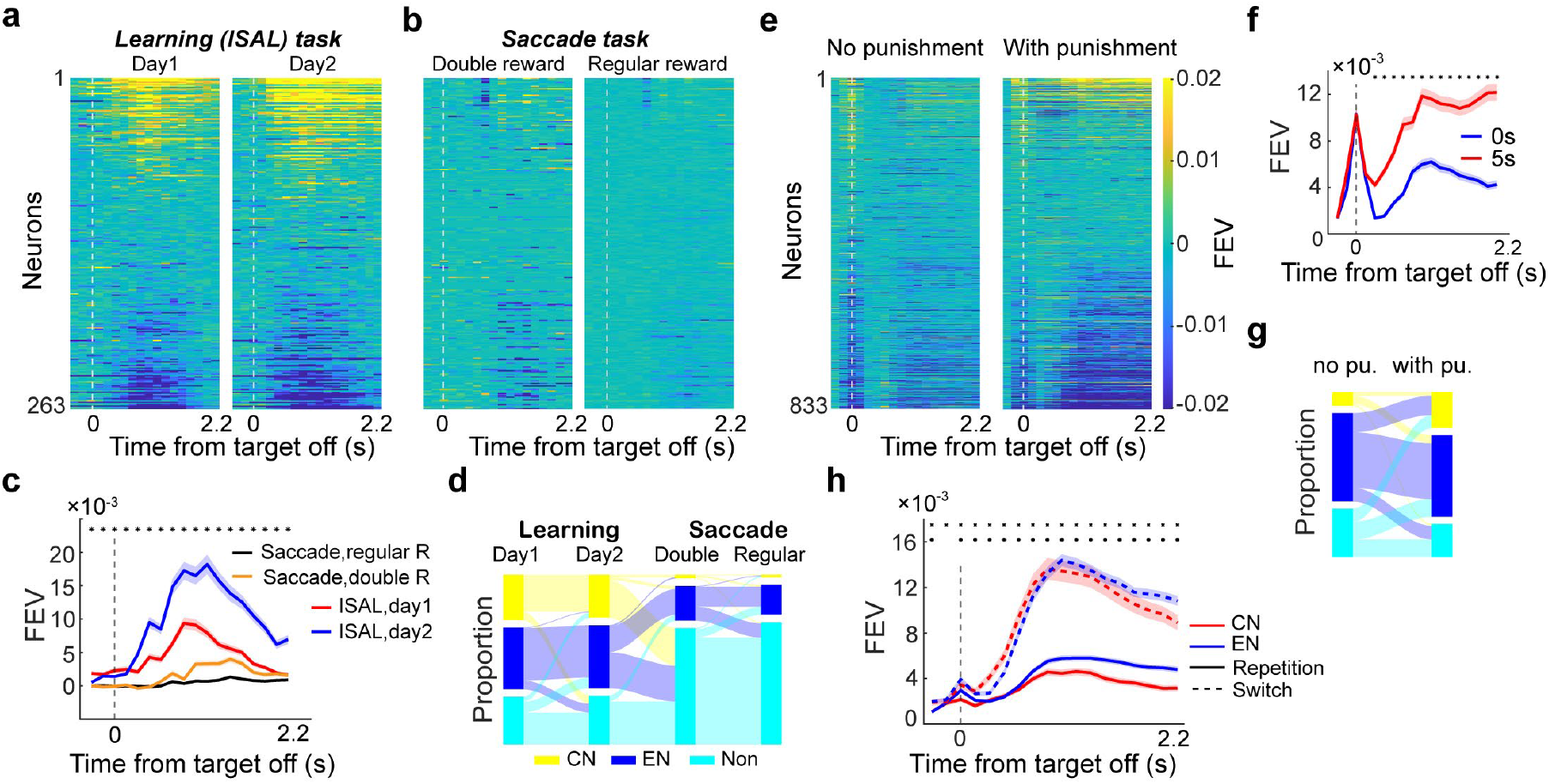
Outcome encoding in 7a did not merely reflect reward reception. **a-d**. Comparison of outcome encodings between ISAL task and a visual saccade task. **a**. The outcome selectivity of 263 7a neurons were shown for two ISAL sessions. **b**. Outcome selectivity of the same neurons were shown for the saccade task, with separate quantification performed for trials using double-sized rewards and trials using regular-sized rewards. Neurons were presented in the same order as in (A) and (B). **c**. Comparison of the averaged outcome selectivity between two tasks. Shaded region denotes SEM. The black stars denote the time points with significant difference between the two tasks (P < 0.01, repeated ANOVA). **d**. Changes in neuron types between different tasks. **e-g**. Comparison of outcome selectivity between sessions using different levels of punishment. **e**. The outcome selectivity of 833 7a neurons were shown separately for the data sessions with and without the time-out punishment. **f**. Comparison of the averaged outcome selectivity between the data sessions with and without the time-out punishment. (P < 0.01, paired t-test). **g**. Changes in cell types between the two task conditions. **h**. Comparison of average outcome selectivity between trials following outcome switches (correct→error or error→correct) and those following outcome repetitions (correct→correct or error→error).

Second, we examined whether outcome encoding in 7a was modulated by punishment. During some of the ISAL recording sessions, we removed the time-out punishment after incorrect trials. This manipulation significantly attenuated the strength of outcome encoding in 7a neurons (**Figure 3e-f**, P = 7.7e-23, t(832) = -10.13, paired t-test). Notably, the deletion of punishment changed the distribution of outcome-selective neurons: the proportion of CNs diminished significantly whereas that of ENs increased notably (**Figure 3g**, P_(EN)_ = 0.011, P_(CN)_ =1.4e-8, chi-square test). Some ENs (12.73%) even switched to encode correctness upon adding punishment.

Third, we investigated whether outcome encoding in area 7a was modulated by outcome history, specifically the outcome of the preceding trial. We quantified the encoding strength separately for trials following outcome switches (correct→error or error→correct transitions) versus outcome repetitions (correct→correct or error→error). Notably, outcome encoding was much stronger following outcome switches compared to outcome repetitions (**Figure 3h**, P = 6.7e-14, t(116) = - 8.5, paired t-test). This effect was consistent across both error-preferring and correct-preferring neurons for both monkeys (**Supplementary Fig. 13**). Together, these results demonstrate that outcome encoding in area 7a extends beyond simple reward reception. It reflects the evaluation of both positive and negative choice consequences, and dynamically incorporates recent behavioral history. These findings align with the established role of the primate posterior parietal cortex (PPC) in salience representation, suggesting that area 7a preferentially encodes behaviorally salient outcomes during AL.

Next, we investigated the relationship between outcome encoding in area 7a and the monkeys’ learning behavior during AL. If outcome encoding in 7a supports outcome monitoring and contributes to sensorimotor AL, we would expect that the strength of population-level outcome encoding correlates with the monkeys’ choice strategies and behavioral performance over the course of learning. To test this, we first assessed whether outcome encoding strength was associated with the extent to which monkeys relied on an outcome-guided strategy (win–stay or lose–shift) versus an outcome-ignored strategy (win–shift or lose–stay) during learning. We modeled the monkeys’ saccade choices for each image-saccade association using the PsyTrack model, which estimates a trial-by-trial weight for outcome-guided strategies. Across different learning days, we found a significant positive correlation between the outcome-guided strategy weight and the strength of outcome encoding in area 7a (**Figure 4a-b**; Pearson’s r = 0.54; Spearman’s r = 0.49). This result indicates that outcome encoding in 7a tracked the monkeys’ use of outcome-driven strategies, supporting its role in trial-by-trial outcome monitoring during AL.

**Figure 4.**
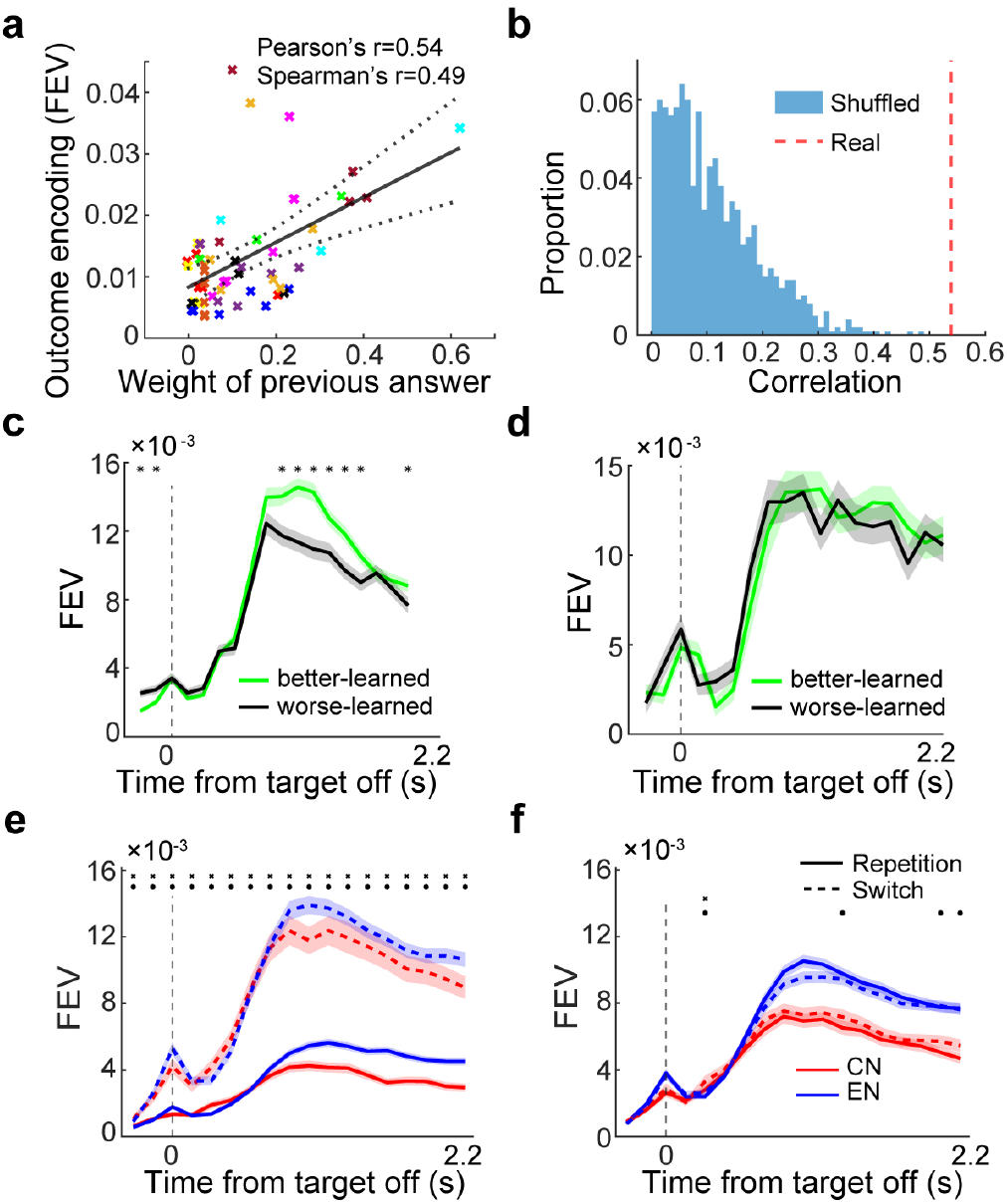
outcome encoding in area 7a correlated with monkeys’ learning behavior. **a**. Correlation between the strength of outcome encoding and the weight of previous answer (extent to which monkeys adopted an outcome-guided strategy, i.e., win–stay or lose–shift) during learning. Each cross represents one recording day, and different colors represent different FOVs. **b**. Histogram of the Pearson correlation values calculated by shuffling the learning days. The correlation value calculated from the real data was significantly greater than that from the shuffled data, indicating a meaningful relationship. **c-d**. Comparison of the averaged outcome selectivity between the better-learned and worse-learned associations. **c**. Sessions with a significant difference in performance accuracy between the two associations. **d**. Sessions with similar levels of performance accuracy. **e**. Comparison of the averaged outcome selectivity between trials preceding exploratory choices (monkeys switched saccade direction relative to the previous trial for the same image–saccade association) and trials preceding exploitative choices (monkeys repeated the same saccade direction for the same association). **f**. Comparison of the averaged outcome selectivity between trials preceding saccade switches and saccade repetitions, regardless of the image–saccade association. The black dot denotes the time point where the outcome encoding was significantly stronger in trials prior to switching choice than repetition choice.

Second, we examined whether outcome encoding in area 7a was associated with the monkeys’ learning performance. For each learning day, we categorized the two tested associations as either better-learned (higher accuracy) or worse-learned (lower accuracy), based on the monkeys’ performance accuracy. In sessions with a notable performance difference between the two associations, 7a neurons exhibited significantly stronger outcome encoding for the better-learned association compared to the worse-learned one (**Figure 4c**; P = 1.56e-7, t(17) = -8.5, paired t-test), regardless of whether the better-learned association was linked to contralateral or ipsilateral saccades (**Supplementary Fig. 14**). This difference in outcome selectivity is unlikely to be attributable to saccade direction encoding alone, as saccade choices were made at least one second prior to target offset. In contrast, in sessions where behavioral performance was similar between the two associations, outcome encoding remained comparable across conditions (**Figure 4d**; P = 0.68, t(8) = 0.42, paired t-test). Together, these results indicate that outcome encoding in area 7a is closely linked to both the monkeys’ behavioral strategies and their learning performance, reinforcing its proposed role in outcome monitoring during AL.

Furthermore, we investigated whether outcome selectivity in area 7a directly influenced the monkeys’ learning of associations. Trials were categorized into two conditions: exploration, where monkeys switched saccade direction relative to the previous trial for the same image–saccade association, and exploitation, where they repeated the same saccade direction. For each condition, we assessed outcome selectivity in the preceding trial. Outcome selectivity was much stronger prior to exploratory trials than exploitative ones (**Figure 4e**, P = 6.94e-13, t(115) = -8.1, paired t-test), suggesting that enhanced outcome encoding in area 7a predicted subsequent exploratory behavior. This effect was observed in both correct-preferring and error-preferring neurons across both monkeys (**Supplementary Fig. 15a-b**). In contrast, 7a neurons did not exhibit elevated outcome selectivity prior to trials where monkeys switched saccade direction independently of the image–saccade association (**Figure 4f** and **Supplementary Fig. 15c-d**). These findings suggest that outcome representations in area 7a may contribute specifically to strategy shifts during learning sensorimotor associations, rather than reflecting general motor variability. Additionally, we found that pre-saccadic selectivity of 7a neurons was significantly, though very modestly, stronger when the monkey’s saccade choice was correct in the preceding trial compared to when it was incorrect (**Supplementary Fig. 16**), suggesting that outcome signals may modulate the strength of association encoding in area 7a in the subsequent trial. Together, these findings indicate that outcome encoding in area 7a fulfills all three proposed criteria for outcome monitoring, suggesting that this region plays an active role in monitoring trial outcomes and contributes functionally to sensorimotor AL.

### Evolution of outcome representation during associative learning

Building on the above findings that establish a correlation between outcome encoding in 7a and outcome monitoring during AL, we next delved into the dynamics of outcome encoding across different learning days during long-term AL. We tracked the activity of neurons in 12 FOVs spanning the sensorimotor AL course (**Supplementary Fig. 17** and **Supplementary Table 1**). The longitudinal recordings for each FOV spanned from the initial day when monkeys performed the ISAL task with familiar associations to the day when monkeys mastered a new pair of associations (**Figure 1b, Figure 5** and **Supplementary Fig. 17**). We assessed the changes in preference and strength of outcome encoding between familiar days (when monkeys were tested with familiar associations) and learning days (when monkeys were learning novel associations). **Figure 5a-j** illustrates the progression of outcome encoding in neurons from two representative FOVs. For the FOV in **Figure 5a-e**, the number of neurons showing outcome selectivity gradually increased after the monkey transitioned to learn novel associations. Within this example FOV, most neurons (76%), initially devoid of outcome encoding during familiar day, developed significant outcome encoding during learning days; while some of the remaining neurons (13.3%) switched outcome encoding upon acquiring new associations (**Figure 5c-e**). At population level, outcome encoding in this FOV progressively increased during the learning days (**Figure 5d**). Conversely, the other FOV exhibited a distinct evolution of outcome selectivity (**Figure 5f-j**). Following an initial surge on the first learning day, outcome selectivity gradually waned over subsequent learning days. This decline is evident from the diminishing outcome selectivity in individual neurons and a reduction in the number of neurons significantly selective for outcomes (**Figure 5h-j**). We also identified FOVs in which outcome selectivity was strongest during the familiar day, and dramatically diminished after transitioned to learn novel associations (**Supplementary Fig. 18**).

**Figure 5.**
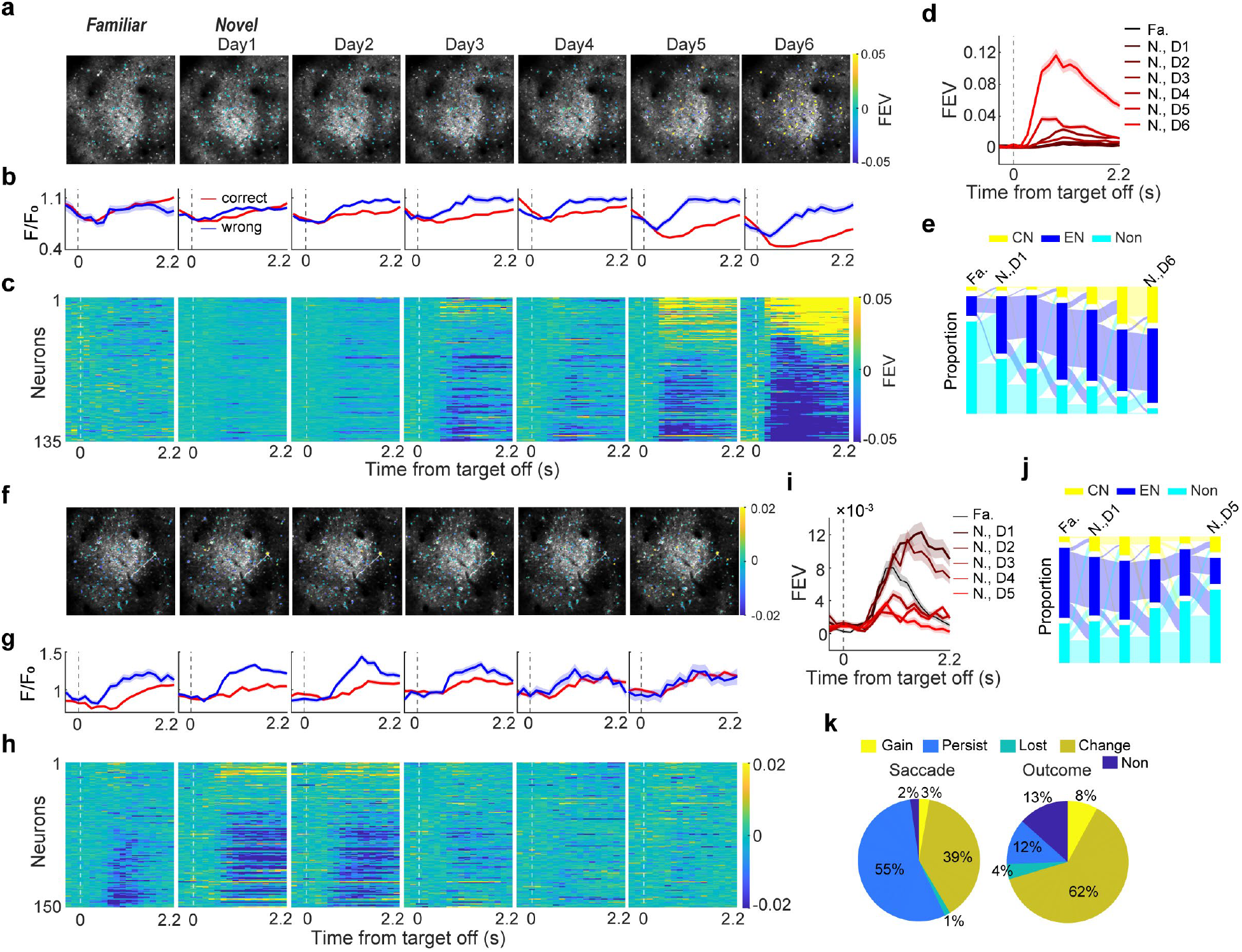
Evolution of outcome encoding in 7a during AL. **a-e**. Evolution of outcome selectivity within one example FOV during an example learning course. **a**. The average two-photon images of this FOV for different days. **b**. An example neuron in this FOV. **c**. Outcome selectivity of all neurons identified within this example FOV were shown for different days within the learning course. Neurons in different sessions\days were listed in the same order. **d**. The averaged outcome selectivity of all neurons identified in this example FOV. **e**. Changes in cell types in the example FOV across different days. **f-j**. Evolution of outcome selectivity in another example FOV during a separate learning course. **k**. Proportion of neurons that changed selectivity over different days during AL. The outcome selectivity and saccade direction selectivity were quantified separately. Data from all 12 learning courses were pooled together. Fa.: familiar day; N., D1: novel stimuli, day 1; N., D2: novel stimuli, day2; and so on.

We found changes in outcome selectivity during AL in most of the tested FOVs (**Supplementary Fig. 19**), although the patterns of change were not unitary. Across all the FOVs, 12% of neurons gained or lost outcome selectivity after monkeys learned novel associations (**Figure 5k**), 62% of neurons changed their outcome selectivity during the learning course (decrease, increase, or switch encoding preference, see Methods), while only 12% of neurons maintained stable outcome encoding across the learning course. Notably, changes in outcome encoding were more prevalent than the changes in saccade direction encoding in 7a, as significantly fewer neurons displayed significant changes in post-saccadic selectivity over the learning course (**Figure 5k**, p << 0.001, chi-square test).

Subsequently, we investigated the stability and evolving patterns of outcome encoding at the population level throughout each learning course through two distinct approaches. First, we calculated the correlations in the patterns of outcome selectivity for all identified neurons in each FOV between successive days (see Methods). This revealed generally low correlation values for most FOVs (r = 0.345±0.146). The inter-day correlations found for outcome encoding were significantly lower than those for saccade direction encoding (**Figure 6a**, P =3.4e-4, t (11) =-5.12, paired-t test), indicating that outcome encoding changed faster than the saccade direction encoding during AL. The patterns of neural encodings on trial outcome and saccade direction were distinct, with modest population-level correlation in only 1 of 12 FOVs (r = 0.015, P = 0.008). Importantly, we found lower inter-day correlations for outcome selectivity between the familiar day and the first learning day than that between successive learning days (**Figure 6a**, P = 0.035, t(11) = -2.24, paired t-test). Conversely, neural encoding of saccade direction did not exhibit such divergent patterns of changes across different learning periods. This suggests that the neural pattern of outcome encoding reorganized in 7a when monkeys began to learn novel associations. The remapping of outcome encoding was notable after learning new associations, as evidenced by significantly lower correlations of outcome encoding between the familiar day and the final learning day (when the monkey had learned a novel pair of associations), compared to the inter-day correlations between successive learning days (**Figure 6a**).

**Figure 6.**
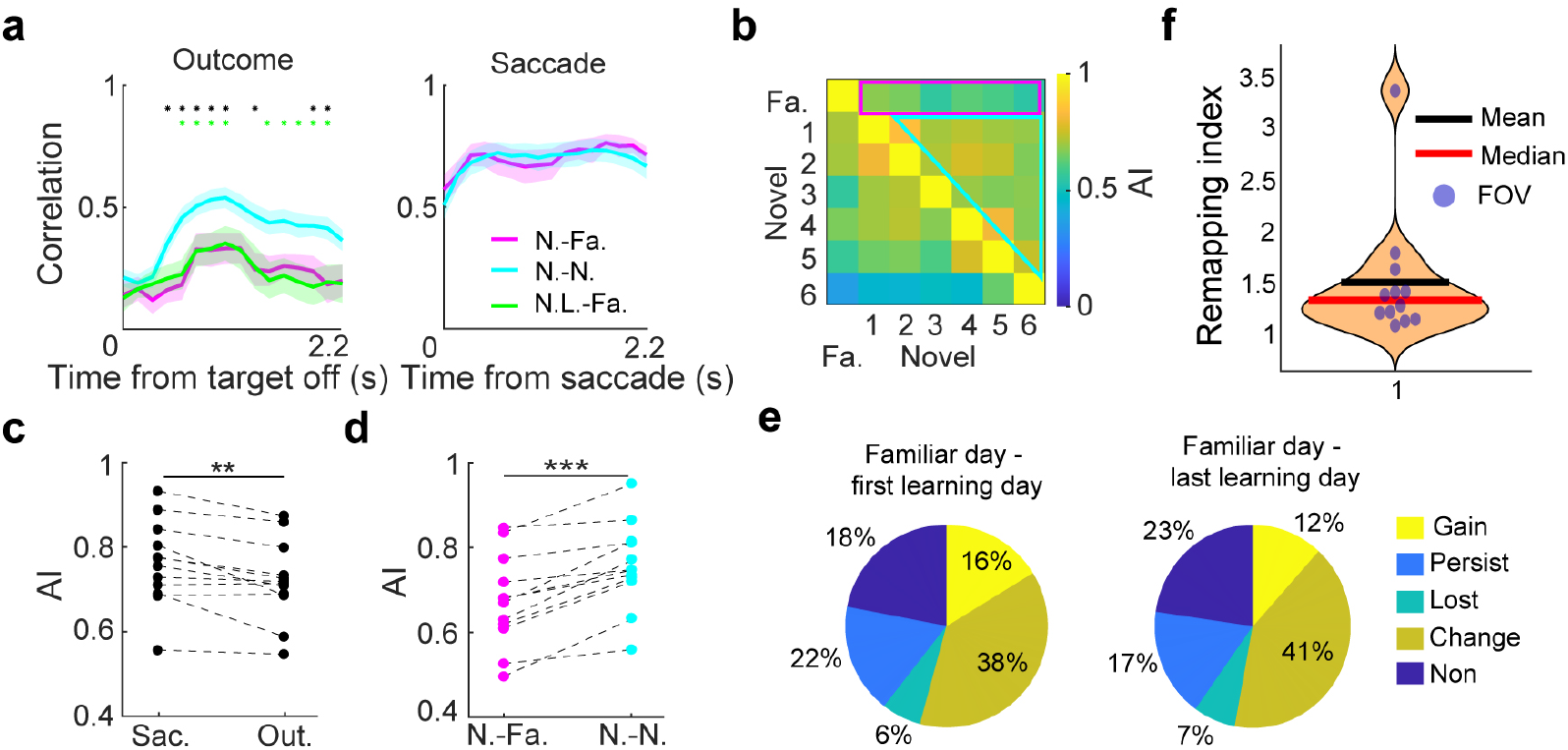
The neural representation of trial outcome reorganized after monkeys transitioned to learning novel associations. **a**. Correlations of neural encodings between two consecutive days (see Methods), averaged for all 12 FOVs. Correlations between the familiar day and the first learning day (N.-Fa., purple) and correlations between different learning days (N.-N., cyan) were compared. The correlation of outcome encodings between the familiar day and the last learning day (N.L.-Fa.) were shown in green. Shaded region denotes SEM. Stars indicate the time points where a significant difference was observed between the N.-N. condition and the other conditions. **b**. Illustration of alignment index (AI) calculation for one example FOV during a learning course. The averaged AI value resulting from the projection of population activity on learning days onto the top principal components (PCs) of neural activity on familiar day (marked by the pink rectangle) quantifies population activity similarity between familiar and learning days (N.-Fa.). Similarly, the averaged AI value resulting from the projection of population activity on later learning days onto top PCs of neural activity on earlier learning days (marked by the cyan triangle) quantifies population activity similarity between different learning days (N.-N.). **c**. Comparison of AI values between saccade direction encoding and outcome encoding. Each data point denotes the averaged result for one FOV. **d**. Comparison of AI values for outcome encoding between N.-Fa. and N.-N. conditions. **e**. Proportion of neurons that changed selectivity after monkeys transitioned to learning novel associations. The results from comparisons between the familiar day and the first learning day, as well as between the familiar day and the last learning day, are shown separately. Data from all 12 learning courses were pooled. **f**. Remapping index quantifying the extent of outcome encoding remapping caused by learning novel associations. Each data point represents the result of one FOV.

Second, we investigated how the population neural codes for trial outcome changed throughout learning. Specifically, we quantified the degree of alignment (or overlap) between the neural subspaces of the population activity from different days within each learning course, and tested how the neural subspaces encoding trial outcome or saccade direction changed throughout the learning course. This approach compares the percentage of variance explained when projecting population activity from one day onto neural axes defined by population activity from another day, producing an alignment index (AI) ranging from 0 (perfect orthogonality between the two subspaces) and 1 (perfect alignment; **Figure 6b** and **Supplementary Fig. 20;** see Methods). We found significantly lower AIs for trial outcome compared to saccade direction in most FOVs (**Figure 6c**), indicating faster changes in neural encoding of trial outcome than that of saccade direction. We then split the AIs into two groups (**Figure 6b**): 1) AIs from projecting neural activity on learning days onto neural axes defined by neural activity from familiar days (novel-familiar group); and 2) AIs calculated using neural activity in different learning days (novel-novel group). Remarkably, AIs for outcome encoding in the novel-familiar group were significantly smaller than that in the novel-novel groups (**Figure 6d**). This difference resulted from neither the difference in monkeys’ behavior performance nor the difference in the temporal gaps between familiar day and learning days (**Supplementary Fig. 21a-b**). Furthermore, the AIs from projecting neural activity on the final learning day onto neural axes defined by activity from familiar days were significantly lower compared to those in the novel-novel condition. Together, these results demonstrate that outcome encoding in 7a was contingent on learning content, and evolved gradually during learning novel associations.

In order to excess the extent of outcome encoding remapping caused by learning novel associations, we further calculated the percentage of neurons whose outcome selectivity significantly changed after the monkeys transitioned to learning novel associations. For each learning course, we compared the outcome encoding on the first or last learning day with the outcome encoding on the familiar day. We found that the majority of neurons (60%) exhibited significant changes in outcome selectivity after transitioning to learn novel associations (**Figure 6e**), while only a small proportion of neurons (17%-18%) maintained stable outcome encoding. To further quantify the extent of such reorganization in population-level outcome encoding, we defined a remapping index as the ratio of inter-day change in population-level outcome encoding (measured by the alignment index) for the novel-familiar group compared to the novel-novel group (see Methods). A higher remapping index indicates a greater degree of change in outcome encoding resulting from the transition to novel associations, relative to the evolution of outcome encoding during the learning of the same associations. Values greater than 1 suggest a trend of remapping, with higher values reflecting more significant changes. We found that the remapping index for all the FOVs was greater than 1, with an average value close to 1.5 (**Figure 6f**), suggest that the reorganization of outcome encoding due to the transition to novel associations was substantial.

Moreover, changes in outcome encoding during AL were not primarily due to variations in the signal to noise ratio of the recorded neurons across different days. First, AIs for outcome encoding calculated using data from two successive days showed no significant difference from the AIs calculated using half-session data within each day (**Supplementary Fig. 21c**). Second, changes in outcome encoding across different days did not correlate with changes in baseline activity of each recorded neuron (**Supplementary Fig. 21d**).

### Network connectivity constrained the evolution of outcome encoding

We hypothesized that the evolution of outcome representations during AL is constrained by network structure. One functional measure of network connectivity is ‘noise correlation’^49,50^, the covariation in activity across repeated trials. We calculated the noise correlation between each pair of recorded neurons for each FOV. This reveled that noise correlation was greater for within-type pairs than cross-type pairs for the outcome-selective neurons (ENs /CNs, **Figure 7a**), indicating that neurons that showed similar neural encodings were more likely to coactivate. We therefore tested whether the strength of noise correlation could predict the changes of outcome selectivity across different days during AL. If the evolution of outcome encodings in 7a was governed by the network connectivity, we expected that neurons that coactivated more closely (more noise correlation) on one (henceforth ‘current’) day would demonstrate similar neural encodings (higher signal correlations) on the next day. Indeed, we found significant correlations between the noise correlation patterns on the current day and the signal correlation patterns on the next day for all successive AL days. (**Figure 7b**, maximum P = 6.9e-64). Additionally, changes in signal correlations were significantly correlated with changes in noise correlations for all successive learning days (**Figure 7c**, maximum P = 5.5e-19). These correlations suggest that the network connectivity potentially constrained the changes in the outcome encoding during long-term AL.

**Figure 7.**
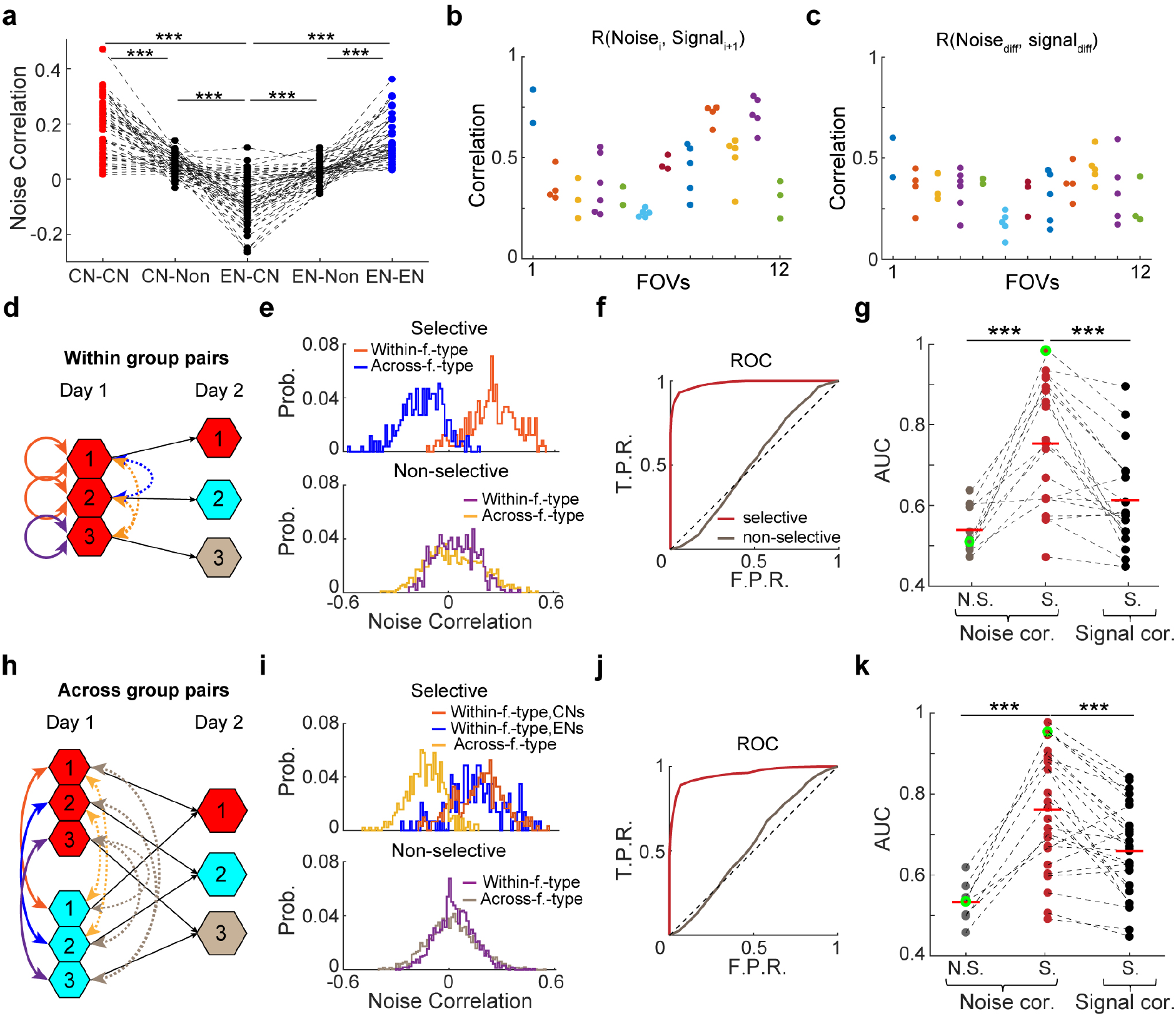
Network connectivity governed the evolution of outcome encodings during AL. **a**. Comparisons of the noise correlation between different neuron pairs. Each data point denotes the averaged value of noise correlation across one group of neuron pairs recorded from one session. **b**. The Pearson correlations between noise correlations of 7a neurons in the current day and the signal correlations of these neurons in the subsequent day. Different colors denote different learning courses. Each dot denotes the result from two consecutive days. **c**. The correlation between the changes in noise correlations and the changes in signal correlations of 7a neurons across two consecutive days. **d**. Schematic illustration of noise correlation calculation for the within-type pairs. This analysis only included neurons that belonged to the same type (CN, EN, or non-selective) on the current day but transitioned to different types on the next day. In day 2, neuron type 1 and type 2 are outcome-selective (S.), while neuron type 3 is non-selective (N.S.). Colored solid and dashed arrows indicate the within-future-type pairs and across-future-type pairs, respectively. **e**. Noise correlation distributions for an example session. **f**. An ROC analysis was used for quantify the difference in noise correlation between within-future-type pairs and across-future-type pairs. The area under curve (AUC) value quantifies the predictive power of noise correlation for neuron ensemble evolution. The red and gray lines indicate the ROC curves for outcome-selective and outcome-non-selective neurons, respectively. Data are from the same example session shown in (E). **g**. Comparison of the predictive power for neuron ensemble evolution between noise correlation and signal correlation. Each dashed line connects data from the same day. Red and gray dots represent results from outcome-selective and outcome-non-selective neurons, respectively. The red horizontal line represents the average value. The green dots denote data from the example session. Noise correlation was significantly different between within-future-type and across-future-type pairs for neurons which were outcome-selective on the second day, but no such difference was observed for neurons which were outcome-non-selective on the second day. **h**. Schematic illustration noise correlation calculation the for the across-type pairs. This analysis only included neurons that were initially of different types on the current day but merged to the same type on the following day. **i-j**. Difference in noise correlations between within-future-type pairs and across-future-type pairs in another example session. **k**. Comparison of the predictive power for neuron ensemble evolution between noise correlation and signal correlation. (***: p<0.001, paired t-test). Data in (j), (k) were shown in the same format as in (f), (g).

To further investigate how the evolution of outcome encoding was governed by the local network connectivity, we examined whether the noise correlation could predict changes in neuron types during AL. We focused on two major groups of neurons whose outcome encodings changed across two concessive days. First, we tested neurons belonging to the same type (CN, EN, or non-selective) on the current day but transitioning to different types on the next day. We categorized these neurons into different subgroups based on their outcome-selectivity on the next day (**Figure 7d-g)**, and calculated the noise correlations (within the same neuron type) on the current day for both within-future-type pairs (neurons belonging to the same type on the next day) and cross-future-type pairs (neurons transitioning to the different types on the next day). This revealed that noise correlation was predictive of the changes of neuron types during AL: noise correlation on the current day was significantly higher for within-future-type pairs than cross-future-type pairs (**Figure 7e-g**). This trend was specifically observed in neurons exhibiting significant outcome selectivity (ENs and CNs), but dramatically diminished in neurons that were nonselective on the next day (**Figure 7e-g**, P = 3.1e-5, t(11) = -6.76, paired t-test). Remarkably, the prediction power for the changes in neuron types significantly decreased when signal correlation was quantified instead of noise correlation (**Figure 7g**, P = 7.7e-5, t(19) = 5.01, paired-t test). These results indicate that the functional connectivity between the same type of neurons could predict the changes in outcome encodings across learning days.

Second, we examined neurons that were initially of different types on the current day but merged to the same type on the following day. Similarly, we quantified noise correlations between these neurons (across different neuron types) on the current day and sorted them into within-future-type pairs and cross-future-type pairs, categorized by the outcome selectivity of these neurons on the next day (**Figure 7h-k**). This also disclosed significantly higher noise correlation on the current day for within-future-type pairs than cross-future-type pairs (**Figure 7i-k**), with this pattern evident only among neurons displaying significant outcome selectivity on the next day (**Figure 7i, k**, P = 5.3e-5, t (7) = -8.71, paired-t test). Furthermore, noise correlation exhibited better predictive ability for changes of neuron types than the signal correlation between these neurons (**Figure 7k**, P = 7.1e-5, t(24) = 4.79, paired-t test). These results suggest that neurons exhibiting closer coactivation on the current day were more likely to develop similar neuronal encodings on the next day, even though they were currently of different types. Together, our results indicate that network connectivity shaped the evolution of outcome encoding in 7a during long-term AL.

## Discussion

By long-term tracking of neural activities of the same neuron groups within the primate PPC during sensorimotor AL, we demonstrated robust neural representations of the trial outcome that was closely correlated with monkeys’ learning. Notably, outcome encoding in area 7a primarily reflected outcome switches, and may directly drive the exploratory behavior in the subsequent trial. Furthermore, these neuronal representations of the trial outcome reorganized as monkeys transitioned from exploitation to exploration and underwent a gradual evolution across learning days. This changing dynamic of outcome representations was notably constrained by the local network connectivity. Our study reveals a new function for primate PPC - monitoring the salient trial outcome and driving exploration during reward-based learning. While previous studies emphasized fronto-striatal dopaminergic network as the central hubs for reward-based learning^19-22,24^, our findings extend the scope of this network and broadens our understanding of the function of the PPC beyond its traditionally recognized role in sensorimotor processing. Additionally, our results reveal for the first time the evolution of the neuronal representation in the same population of neurons in the primate brain during the learning process over a long period, along with the potential mechanisms that govern the evolution of neural selectivity within local networks during learning.

The outcome signal observed in area 7a exhibited several notable features that support its important role in outcome monitoring during sensorimotor AL. First, the outcome encoding in 7a dramatically diminished in the simple saccade task that involved no learning. Second, the strength of outcome encoding significantly decreased when the time-out punishment was removed for the incorrect trials. These results suggest that the outcome representation in 7a transcends simple physical reinforcement feedback, implying involvement of cognitive processing. Third, the strength of outcome encoding in area 7a correlated with both the extent to which the monkeys’ saccade choices followed a win-stay, lose-shift strategy and their performance accuracy during AL. Fourth, the outcome encoding in 7a was contingent on the specific learning content and gradually evolved during AL. This strong reliance of outcome representation on monkeys’ learning behavior implies a functional contribution of the outcome signal in 7a to AL. Fifth, outcome representations in area 7a were particularly sensitive to outcome switches. This aligns with the established role of the primate PPC in salience representation^32,33^. Finally, stronger outcome encoding in area 7a predicted exploratory choices in the following trial but did not predict improved learning performance. This indicates that while outcome signals in 7a may not directly facilitate the formation of sensorimotor associations, they may instead promote exploratory behavior during the learning process. Together, these results suggest that the primate PPC satisfy the three criteria for outcome monitoring. It contributes to sensorimotor AL by monitoring behavioral outcomes to facilitate exploratory behavior, extending its recognized role in reward-based decision-making.

Previous studies have identified outcome representation in brain areas outside PPC, emphasizing the significance of the frontal-striatum network in evaluating reward reinforcement^3,19-21^. Outcome signal has been proposed to serves as a teaching signal, reinforcing correct associations while weakening incorrect ones. However, the observed outcome representation in PPC differs markedly from these earlier findings, instead highlighting a predominant sensitivity to salient, negative outcomes that may drive exploratory choices during AL. This suggests a potential complementary role of PPC in monitoring trial outcomes during sensorimotor AL, relative to the role of the frontal-striatum network. While the frontal-striatum circuitry primarily evaluates reward reinforcement and supports the acquisition and exploitation of learned associations during AL, PPC may instead play a predominant role in detecting salient ‘error signals’ and facilitating exploratory behavior throughout the learning process. The outcome signal in area 7a may be primarily relayed to brain regions such as the frontal pole and intraparietal sulcus, which are known to drive exploratory behavior during decision-making^51,52^. This is also in line with opponent-process theories, which propose that emotional experiences and motivated behavior are regulated by pairs of opposing processes^53^.

The critical roles of the fronto-striatal network in AL have been demonstrated in previous studies using causal approaches, such lesion studies^24,54-57^. However, the causal involvement of the primate PPC in long-term AL remains to be tested in future studies. Our findings suggest that area 7a may not play a central role in forming or encoding sensorimotor associations during AL. First, only a small fraction of 7a neurons showed significant encoding of the visual cue during either the cue presentation (<4%) or delay (20%) periods of the ISA task. Second, encoding of saccade direction prior to the monkeys’ choices was notably weak. Third, this pre-saccadic encoding did not show systematic changes across learning days that aligned with the monkeys’ learning performance. The relative lack of attention to the primate PPC in earlier AL studies aligns with our observation that area 7a may not be directly involved in acquiring new associations. Instead, it may contribute to outcome monitoring—specifically by encoding salient outcome signals that promote exploratory behavior during AL. Nonetheless, the role of the primate PPC in outcome monitoring and its potential contribution to exploration-driven learning processes warranting further investigation with causal manipulations. Previous rodent studies have reported that PPC was causally involved in processing sensory-stimulus history and history-based action selection bias to guide decision behavior^58,59^. Moreover, PPC was also shown to represent discrepancies between the current model and observed state transitions in reinforcement learning^60^, as well as reward-related belief updating during flexible decision-making^61^. Collectively, these results indicate that PPC likely plays an important role in updating the estimation of the current state by incorporating past experience, to guide decision-making and learning.

Previous studies have shown that neural activity in area 7a reflects saccade direction^62,63^, eye position-induced gain fields^64^, and reaching movements^65,66^. However, several lines of evidence suggest that the outcome encoding observed in our study is unlikely to originate primarily from sensorimotor processes. First, the outcome encoding cannot be attributed to differences in the monkeys’ eye positions or eye movements between correct and error trials during the inter-trial interval. Second, outcome encoding in area 7a significantly diminished when the task did not involve learning or when the time-out punishment for incorrect trials was removed. Third, outcome encoding underwent substantial reorganization when the monkeys transitioned to learning novel associations and then gradually evolved in distinct patterns during the learning of different novel associations. Nevertheless, it is possible that the monkeys exhibited idiosyncratic but systematic behavioral changes during the inter-trial interval, such as limb movements, sucking, or body gestures, while learning different novel associations and engaging in various task conditions. Furthermore, it is also possible that the monkeys were engaged differently across tasks or learning periods, which might be reflected by changes in pupil signals. These behavioral changes could partially contribute to the outcome encoding in area 7a. Future studies should further investigate the relationship between these idiosyncratic behaviors and outcome representation during the learning process.

Remarkably, our investigation revealed that the outcome encoding in 7a was contingent on the learning content. The neuron ensembles that encoded the trial outcome substantially reorganized when monkeys transitioned to acquiring new association, and the outcome encoding of individual 7a neurons gradually evolved over days during learning. This episodic-like outcome representation, however, did not signify a global remapping of neural representations in 7a after transitioning to new associations, nor did it primarily stem from the overall fluctuation of neural encoding across different days, because such significant reorganization and rapid changes were not observed for the representation of saccade directions. Consistently, stronger outcome selectivity was specifically linked to switching saccade choice direction for the same imaging-association rather than simply switching saccade directions irrespective to the associations. This content-dependent outcome encoding may also explain the heterogeneous evolution of outcome signals across different 7a subregions (FOVs) throughout the learning process. Beyond potential clustering of outcome-encoding neurons, such heterogeneity might reflect the storage of distinct learning experiences or support the acquisition of different types of content.

Consistently, previous rodent studies have shown that the firing patterns of grid cell and space cells remapped after transitioning to new environment^67^. This remapping process is thought to incorporate information about space together with particular events or function as a pattern separation mechanism, both of which are essential for episodic memory processing^68^. Similarly, the reorganization of outcome representation in PPC might incorporate the outcome information with distinct sensorimotor associations during AL, potentially segregated in memory as discrete learning experiences. Consequently, new patterns of outcome representations emerged as monkeys embark on acquiring new associations. We speculate that PPC neurons exhibiting strong outcome encoding during the learning of specific associations may become functionally coupled with neurons representing these associations, thereby influencing the learning of sensorimotor associations. Such content-dependent outcome representations may provide a computational advantage over abstract, content-independent outcome signals by enabling outcome-encoding neurons to directly influence the neuronal ensembles representing specific associations, without requiring more complex targeting mechanisms. Further studies—such as reversing the sensorimotor associations and causally manipulating activity dynamics—are needed to clarify the potential role of encoding remapping during AL.

Our study had explored the neural mechanisms underlying long-term learning in behavioral monkeys. Earlier work has demonstrated that long-term training shapes neural activity in the primate brain^25-27^. Our work extends these results not only in demonstrating how AL shapes the representation of individual neurons across different learning days, but also by providing insights into the mechanisms that constrain the evolution of neural selectivity during learning. Employing noise correlation to measure the functional connectivity between single neurons, we found that the changes in neuron ensembles was casually constrained by the network connectivity. While noise correlation is commonly used to quantify functional connectivity between neurons, previous studies have primarily focused on its impact on population-level sensory encoding or how it changes during learning^25,50,69-71^. Our results indicate that the structure of correlated activity variability may also be useful for exploring learning-related plasticity of cortical connectivity. Specifically, our results indicate that neurons coactivated more closely were more likely to develop strong connections over time, leading to similar neural encoding. This aligns with the Hebbian theory, which posits that neurons that fire together wire together. Furthermore, our work demonstrated that noise correlation among neurons could better predict changes in neuronal encoding after learning as compared to the similarity of their functional properties. Together, these results suggest a potential foundational principle for learning: the network structure serves as a constraint on learning processes.

## Acknowledgments

We thank Prof. Muming Poo, Prof. John Maunsell, Prof. David J. Freedman, Prof. Samuel Solomon, Prof. Tianming Yang, Prof. Lusha Zhu for their helpful comments on an earlier version of this manuscript. We also thank the veterinary staff at Peking University Animal Resources Center for expert assistance.

## Funding

Ministry of Science and Technology of the People’s Republic of China, STI2030-Major Projects (2021ZD0203800), YZ;

National Natural Science Foundation of China, NSFC32171036, YZ.

## Author contributions

Conceptualization: YZ

Methodology: ZL, LS, ST

Investigation: ZL, LS, ZJ, YZ

Visualization: ZL, ZJ, YZ

Funding acquisition: YZ

Project administration: YZ

Supervision: YZ

Writing – original draft: YZ

Writing – review & editing: FF, YZ

## Competing interests

Authors declare that they have no competing interests.

## Data and materials availability

All data and analyses necessary to understand and assess the conclusions of the manuscript are presented in the main text and in the supplementary materials.

The online link to the raw data and analysis codes will be provided upon acceptance of the manuscript.

## Supplementary Information

**Supplementary Fig**. 1-21

## Methods

### Animal preparation and Behavioral Training

We trained two male Rhesus Macaques, aged 4-5 years and weighing 5-6 kg, on two different behavioral tasks. Before behavior training, we implanted three head-posts in an inverted Y-shape configuration on the monkeys’ skulls. A T-shaped steel frame was affixed to these head-posts to enhance the stability of the head fixing. Our surgical and behavioral approaches have been described in detail previously^72,73^. The monkeys were individually housed in cages, subjected to a 12-hour light/dark cycle. Behavioral training and experimental recordings were exclusively conducted during the light phase. Monkeys were comfortably seated and head-fixed in a custom-made primate chair inside a dark experiment rig. Task stimuli were displayed on a 27-inch color LCD monitor (1920*1080 resolution, 75 Hz refresh rate, 45 cm viewing distance). A solenoid-operated reward system was used to deliver juice reward to the monkeys. Monkeys’ eye positions were monitored by an optical eye tracker (ISCAN ETL-200) at a sampling rate of 240 Hz and stored for offline analysis. Stimulus presentation, task events, rewards, and behavioral data acquisition were accomplished using an Intel-based PC equipped with MonkeyLogic software running in MATLAB^74^ (http://www.monkeylogic.net). All experimental and surgical procedures were in accordance with Peking University Animal Care and Use Committee.

### Behavioral tasks

#### Image saccade association learning task (ISAL)

Monkeys had to learn the associations between different visual images and two saccade directions in this task. On each trial, monkeys were required to saccade to one of the two targets based on a presented visual image (**Figure. 1a**). Each trial started with a central fixation point, and monkeys needed to maintain their gaze within a 3° radius of this point until it disappeared. 1500ms after monkeys acquired fixation, a visual image, randomly selected from the internet and characterized by full contrast and a 6.5° diameter, appeared on the screen for 500ms. Importantly, the image was always presented on the contralateral side of the screen relative to the recorded brain hemisphere (7° eccentricities, orientated at 135°). Following a 1s delay, the fixation point disappeared and two saccade targets (white, 0.5° diameter) appeared simultaneously on the left and right side of the screen, equidistant from the center at 7° eccentricities. These targets were positioned orthogonally to the visual image’s orientation (oriented at 45°). The spatial configurations of task stimuli remained constant throughout the study. Monkeys had to saccade to the direction associated with the presented visual image within a 400ms time window, and maintain fixation on the chosen target for 1s to get juice reward. The juice reward was promptly delivered upon the target’s disappearance. As such, the trial outcome, whether correct or erroneous, became quickly evident, even though explicit feedback regarding the outcome of each trial was absent. Once they had fully learned a pair of image-saccade associations, the monkeys were then instructed to learn two novel associations in subsequent days.

#### Visual saccade task

A visual saccade task was used as a control task to test whether the outcome encoding in PPC solely reflected getting\missing reward. Trials were initiated by the monkeys acquiring central fixation. Following a 1500 ms fixation period, the fixation point disappeared, and a visual target was presented at one of four possible locations, evenly spaced at 90° angular intervals with a 7° eccentricity. Two of the target locations corresponded precisely to those in the ISAL task. Monkeys had to make a single saccade to the target within 400 ms, and fixate at target for 1000 ms in order to receive a juice reward. At the behavioral training phase, monkeys received a standard-size juice reward after completing every trial. However, during neural recording sessions, the reward probability was manipulated: in one-quarter of randomly selected trials, monkeys were rewarded with double the usual amount, while the juice reward was omitted entirely in another quarter of randomly selected trials.

#### Task training

Both monkeys were trained with the visual saccade task before neural recording. However, there were slight differences in the training procedures for the ISAL task between the two monkeys. Monkey H received training in the ISAL task for 61 days but had not fully acquired proficiency in the first pair of image-saccade associations before the commencement of neural recording. Consequently, Monkey H continued to learn these initial associations after the neural recording began, spending almost one month to master them. In contrast, Monkey X received more extended training in the ISAL task and had already fully learned the first pair of associations before neural recording commenced. Both monkeys learned new sets of image-saccade associations more quickly as they gained experience with each new set. Notably, they took significantly longer to learn the first pair of associations, as they had to familiarize themselves with the task rules and stimulus sequence during this initial stage. Over the course of neural recording, two monkeys learned a total of 12 pairs of novel associations (Monkey H: 2 pairs, Monkey X: 10 pairs). This allowed us to consistently track the same neurons across all 12 subregions within 7a throughout the entire learning courses. During each learning course’s neural recording, monkeys were initially tested with the ISAL task using the previously learned (familiar) associations on the first day. Subsequently, they were introduced to a new pair of associations (novel) on the following days until full proficiency was achieved. At this stage of learning, only the novel associations were tested, while the familiar associations were not. Importantly, within each recording session/day, the image-saccade associations remained constant.

While Monkey H was in the process of learning the initial pair of image-saccade associations during the neural recording phase, its performance accuracy stabilized at approximately 80% for a period of several days. During this phase, we removed the time-out punishment following error trials in the ISAL task for four randomly selected sessions/days. This was designed to investigate the impact of punishment on outcome encoding in the PPC. Importantly, all task variables remained consistent within each recording session/day.

### Neural recording

#### Virus injection and chamber implanting

In order to record the activity of a large population of neurons and monitor their activity changes across different days, we implemented two-photon calcium imaging recording on behavioral monkeys. Detailed procedures for the surgical and recording processes are elaborated upon in the previous studies^72^. In brief, we conducted two surgeries on each animal to inject the virus for transferring the calcium indicator into cortex and to implant the optical recording chamber respectively. During the first surgery, a 14-mm diameter craniotomy was executed on the skull over 7a region in the right hemisphere of each monkey. The localization of 7a was determined in accordance with the monkey brain atlas, specifically situated as the posterior part of the inferior parietal gyrus, located beneath the intraparietal sulcus and above the superior temporal sulcus (**Figure 1g-h**). Following the dura mater’s exposure, we performed a pressure injection of 50-100 nL AAV9-Syn-GCaMP6f-WPRE-SV40 (titer 2.6e13 (GC/ml), Addgene) into cortex at a depth of approximately 400 μm. To ensure effective transfection, the virus solution was injected to many sites widely spread in 7a. Approximately 45 days after AAV injection, a second surgery was conducted to implant the recording chamber. A glass coverslip (diameter 8 mm and thickness 0.17 mm) glued in a titanium ring with silicone adhesive (KN-300X, Kanglibang, China) was used to gently cover the cortex. The titanium ring was sealed on the skull with dental acrylic to form an imaging chamber. In both monkeys, tissue regeneration occurred within the imaging chamber several weeks after the chamber implantation surgery. This dramatically degraded the quality of optical signal. We therefore surgically removed the regenerated tissues and replaced the glass coverslip with a new one.

#### Two photon calcium imaging recording

Approximately one week following the implantation of the recording chamber, we initiated the two-photon calcium imaging recording experiment using a Thorlabs two-photon microscope and a Ti:Sapphire laser (Mai Tai eHP, Spectra Physics). In each recording session, a cortical area of 800 μm × 800 μm was imaged at 30Hz using a 16× objective lens (0.8 N.A., Nikon). The recording depth ranged from 200 μm to 300 μm beneath the arachnoid membrane. A standard slow galvo scan was used to obtain static images of neurons with high resolution (1024* 1024). For imaging neuronal activity, we employed a fast resonant scan capable of capturing up to 30 frames per second (averaging each 4 frames resulted in 7.5 frames/s). Each recording session lasted for a duration of 4–6 hours, generating an image data set consisting of 80k-120k frames. Approximately every 5-10 minutes, we manually corrected slow drifts in the field of view (FOV) by comparing them to a reference image. In total, we conducted 117 recording sessions across 25 imaged FOVs distributed in 7a from two monkeys (7 FOVs and 36 recording sessions for Monkey H and 18 FOVs and 81 recording sessions for Monkey X). Within 12 of the 25 FOVs, we consistently monitored the activities of the same neurons throughout a full learning course spanning a period of 3 to 7 days. To ensure alignment with the same imaging plane on consecutive days, we initially performed coarse alignment based on the recording coordinates, blood vessels, pia mater, and arachnoid, followed by precise alignment to reference images.

### Data analysis

#### Behavioral performance

Only recording sessions where the monkeys completed a substantial number of trials (n >= 800 for Monkey H and n >= 700 for Monkey X) were included in our subsequent analyses. On average, monkey H completed 1156 trials per session, and monkey X completed 965 trials per session. We assessed each monkey’s performance accuracy by calculating the ratio of correct trials to the total number of completed trials, encompassing both correct and error trials. In **Figure 1c**, we depicted the real-time accuracy dynamics using a sliding time window approach (width: 100 trials, step size: 20 trials). To determine whether monkeys had fully acquired a pair of image-saccade associations, we applied two criteria: 1) the monkey’s average accuracy throughout the session exceeded 0.8, and 2) the monkey’s average accuracy reached 0.9 over a continuous period of at least 200 trials. Additionally, we assessed whether monkeys exhibited a significant bias in saccade direction within a given session by examining the disparity in their performance accuracies between two different saccade directions. If this performance accuracy difference reached 0.2 within a specific section, we classified that section as a biased session.

##### Dynamic model fitting

In order to capture the dynamics of monkeys’ behavior strategies during sensorimotor AL, we fitted monkeys’ performance accuracy with a dynamic Bernoulli generalized linear model, characterized by a set of smoothly evolving psychophysical weights. Specifically, we used ‘PsyTrack’ package, which was described in detail in a previous study and proven suitable for parametrizing the time-varying behavior in learning task^48^ (https://github.com/nicholas-roy/PsyTrack). This model aims to fit modeling weights that depict the decision-making strategy of the animals on a trial-by-trial basis, as a linear combination of available task variables. Therefore, this model describes choice behavior at the resolution of single trials, allowing us to quantify the evolution of animal’s decision strategies (represented by different weights) across the learning course. Notably, the magnitude of a particular weight directly correlates with the extent to which the animal relies on the corresponding task variable in its decision-making process. Specifically, we utilized the task condition (Cue identity) as input for the model, denoting each single trial as either -1 or 1, while the monkeys’ choice (Saccade direction) served as the model’s output, labeled as 0 or 1 for each single trial. We determined three behavioral weights, namely the bias weight, previous answer weight and stimulus weight, through model fitting.

#### Neuronal data analysis

##### Identify single neurons within each session

We used an image processing software (suite2p, https://github.com/MouseLand/suite2p), for motion correction, definition of putative cell bodies, and extraction of fluorescence traces. We extracted the single neuron activity within each FOV based on three primary steps. First, the raw data streaming from the microscope system was converted into ome.tif format using Fiji software (version 2.13, https://imagej.net/software/fiji/). Subsequently, we transferred this data to ‘suite2p’ for offline processing, which included motion correction, extraction of neuronal signals, and denoising. Second, we manually selected the resulting cell bodies (ROIs) candidates from the original ROI panels. We then employed a machine learning approach to optimize parameters tailored to the data recorded from different subregions of the 7a cortex in each monkey. These refined parameters were subsequently applied to ‘suite2p’ to reprocess the data again, generating new ROIs for each data session. Lastly, we manually inspected and selected ROIs once more to ensure that all identified ROIs exhibited satisfactory neuron morphology and met the quality criteria.

##### Identify the same group of neurons across sessions

Following the selection of ROIs for each recording session, we proceeded to align the neurons across all the recording sessions\days throughout the learning course for each tested 7a FOVs. This involved the following steps. 1) We utilized ‘suite2p’ to generate a cross-day classifier by leveraging data from all sessions/days within each learning course for each 7a subregion. 2) Subsequently, we imported the cross-day classifier back into ‘suite2p’ to generate ROIs within each FOV for each individual day within the learning course. Using these steps, the neurons (ROIs) within each FOV were identified for each recording day through the ‘suite2p’. 3) We then manually cross-referenced and matched the neurons (ROIs) among different recording sessions within each learning course, based on their spatial location in the reference plane and their morphology. If a neuron could not be confidently identified in one of the sessions within the learning course, it was excluded from consideration for all recording sessions within that learning course. Consequently, our approach yielded an incomplete map of all neurons across all recording days for each FOV.

Nonetheless, this method offers advantages over other commonly used approaches. Other approaches often use a single map of ROI masks for all days, such that this map is transformed on each day to best fit that day’s imaging alignment. Slight deviations in the axial plane of the image or other sources of in-plane distortion could lead to slight offsets in masks from day-to-day relative to the ideal alignment. Such slight offsets could result in contamination from activity in other cells, dendrites, and axons. In contrast, our approach identifies signal sources on each day, circumventing the risk of contamination from signal sources on different days. Furthermore, our procedure necessitates the selection of neurons that consistently exhibit a high signal-to-noise ratio throughout the entire learning course. Consequently, we have confidence that the observed changes in neural encoding for each individual neuron during the learning course are unlikely to be attributable to fluctuations in the recording quality across different recording sessions.

##### Neuronal pre-screening, denoise and Normalized activity

Since our recording sessions typically extended for a considerable duration (4-6 hours each), various factors such as slow drifts in the imaging plane’s focus and light bleaching occurred within each session. These factors resulted in gradual alterations in the baseline fluorescence levels of individual neurons during each session. To mitigate these potential sources of interference, we implemented a high-pass filter on the fluorescence trace of each neuron, effectively eliminating very slow dynamics within each recording session (High pass filter, 0.005 Hz). We plotted the relative changes in fluorescence, denoted as F_(t)_/F_0_, for all the examples of single neuron. Here, F_0_ represents the mean activity of the region of interest (ROI) over the entire session, while F_(t)_ signifies the activity of each individual neuron.

#### Quantify the outcome encodings

##### Identify outcome selectivity

To identify neurons with significant encoding of trial outcome, we applied a Wilcoxon ranksum test to assess activity between the correct and error trials in the task period following target offset (1-16 frames (0-2.2s) after target offset). We further divided this period into four smaller windows, each comprising 4 frames. If statistically significant differential activity between correct and error trials was observed within at least one of these windows (P < 0.01, Bonferroni corrected), we classified the neuron as outcome-selective. Similarly, to identify whether the 7a neurons exhibited significant selectivity for the outcome of the previous trial (outcome-history), we compared activity between the trials in which the previous trial was a correct vs. incorrect trial. This comparison included neural activity during the fixation, cue and delay periods of the ISA task (12 frames before to 12 frames after cue onset). We partitioned this time period into six evenly spaced time windows and designated a neuron as outcome-history selective if a statistically significant difference emerged during this comparison (P < 0.01, Bonferroni corrected).

##### Identify saccade direction selectivity

To identify neurons exhibiting significant saccade direction encoding, we employed a Wilcoxon rank-sum test to compare neural activities around saccade onset between trials featuring different saccade directions. Specifically, we assessed saccade direction selectivity during both the pre-saccadic period (1-4 frames before saccade onset) and the post-saccadic phase (1-8 frames after saccade onset). While the pre-saccadic epoch was analyzed as a single time window, we divided the post-saccadic epoch into two smaller time windows (4 frames each). Neurons were classified as pre-or post-saccade selective if there was statistically significant differential activity observed between trials with distinct saccade directions in at least one time window within the pre- or post-saccadic epoch (P < 0.01, Bonferroni corrected).

##### Identify visual responsive neurons

To determine whether 7a neurons exhibited a significant visual response to the visual cue in the ISAL task, we conducted a comparison between each neuron’s activity during cue presentation (1-4 frames after cue onset) and its baseline activity (defined as the fixation period, spanning from 1 to 5 frames before cue onset). Neurons were categorized as ‘visual responsive’ if their activity during cue presentation exceeded their baseline activity by at least one standard deviation.

##### Unbiased fraction of explained variance (FEV)

We performed one-way ANOVA on the neuron’s activity within each time frame to quantify the amount of information that each neuron encoded about trial outcome. Similar as in the previous studies^75,76^, We calculated the unbiased fraction of explained variance (FEV) in the neuron’s activity that could be attributed to the trial outcome with the following:

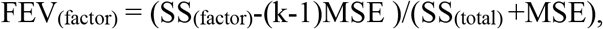

where SS indicates the sum of squares, MSE indicates mean square error, and k indicates number of conditions.

We observed variations in the magnitudes of outcome selectivity across trials in which monkeys executed saccades in different directions. Moreover, monkeys exhibited biased choices toward different saccade directions on different days during AL. Additionally, we recognized that the imbalance in trial numbers between correct and incorrect trials on different days could have varying effects on the FEV value of outcome selectivity. To mitigate potential confounding factors stemming from these phenomena when comparing outcome selectivity across different days, we employed a two-step approach to calculate the FEV value of outcome selectivity for each neuron. 1) Initially, we separately computed the FEV value of outcome selectivity for each of the two saccade directions and subsequently averaged these values. 2) For each saccade direction, we randomly selected an equal number of correct and error trials, matching the smaller of the two trial numbers. We repeated this process 500 times, calculating FEV values each time, and then averaged the resulting 500 FEV values.

##### Determining the latency of outcome selectivity

For each neuron, a Wilcoxon ranksum test was applied to the distributions of activity to the both correct and error trials within each frame to determine the latency of outcome selectivity. The starting point of outcome selectivity was defined as the first of three consecutive frames in which a significant difference (P < 0.01) was observed in neuronal responses between the two trial types.

##### Determining the duration of outcome selectivity

For each neuron displaying outcome selectivity, we divided the task epoch (which was used for identifying outcome selectivity) into multiple time windows, each spanning 4 frames (∼0.54 seconds), to assess the duration of outcome selectivity. Within each time window, we performed a Wilcoxon rank-sum test to compare neuronal activity between correct and error trials. We then determined the number of time windows in which a statistically significant difference (P < 0.05, Bonferroni corrected) was observed.

##### Linear regression model

We examined whether the outcome selectivity of 7a neurons can be simply attributed to encodings relative to differences in monkeys’ eye positions between correct and error trials during the inter-trial interval. Specifically, we employed a multiple linear regression model to quantify the dependence of 7a neuronal activity on monkeys’ eye positions in both horizontal and vertical directions, saccade choice direction, as well as trial outcome:

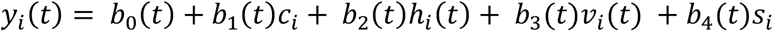

Here, *y*_*i*_(*t*) represent the neuron’s activity at the *t*_th_ frame after the target offset in the *i*_th_ trial, *c*_*i*_ is the trial outcome of trial *i* (1 and -1denotes correct and incorrect, respectively), *h*_*i*_ and *v*_*i*_ represent the horizontal and vertical eye positions relative to the fixation center, and *s*_*i*_ is the saccade choice direction of trial *i*. The coefficients *b*_0_ through *b*_4_ denote the regression coefficients. The linear regression was applied to neuronal activity for each time frame after target offset in the inter-trial interval.

Furthermore, we tested whether the outcome selectivity of 7a neurons was primarily driven by differences in the monkeys’ saccade behavior or blinks between correct and error trials during the inter-trial interval. Saccades and blinks were identified using the following steps: First, we calculated eye movement speed *v*(*t*) during the inter-trial interval after each trial. We marked speed peaks 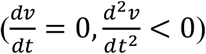 as potential blinks or saccades. Peaks with a maximum speed *v*_*max*_ below 100°/*s* are excluded, whereas those peaks with a speed *v*_*max*_ exceeding 900^°^/*s* are identified as blinks, yielding the blink count or frequency *f*_*b*_ for each trial.

Next, we identified potential saccade starts 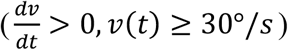 and stops 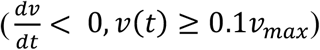 around the remaining peaks. If overlapping saccades were detected, only the one with the higher peak speed was retained. We then verified that all saccades had a magnitude *m*_*s*_ greater than 2° by examining eye position at the start and stop points. Additionally, we checked for consistent saccade direction 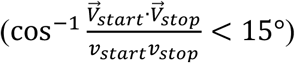 within each saccade. Only movements that passed both criteria were classified as saccades, providing the saccade frequency *f*_*s*_ and saccade magnitudes 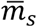 for each trial.

After identifying saccades and blinks during the inter-trial interval, we used the following linear regression model to assess the influence of differences in saccades and blinks between correct and error trials on 7a neural activity:

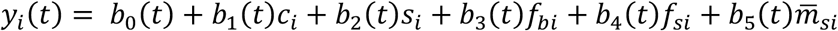

Here, *y*_*i*_(*t*) represent the neuron’s activity at the *t*_th_ frame after the target offset in the *i*_th_ trial, *c*_*i*_ is the trial outcome of trial *i* (1 and -1denotes correct and incorrect, respectively), *s*_*i*_ is the saccade choice direction of trial *i, f*_*bi*_ and *f*_*si*_ represent blink frequency and saccade frequency during the *i*_th_ inter-trial interval, and the 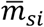 is the average saccade magnitude of those saccades. The coefficients *b*_0_ through *b*_5_ denote the regression coefficients. The linear regression was applied to neuronal activity for each time frame after target offset in the inter-trial interval.

Furthermore, in order to test whether 7a neural activity reflected the outcome history of the previous trials, we employed a multiple linear regression model to quantify the relationship between 7a neuronal activity and the outcome history of previous trials using the following equation:

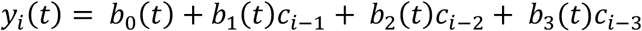

Here, *y*_*i*_(*t*) represents the neuron’s activity at *t*_th_ frame after fixation onset in the *i*_th_ trial, and *c*_*i*_ is the trial outcome of trial *i*, and the coefficients *b*_0_ through *b*_3_ denote the regression coefficients. For this analysis, we focused solely on data from one monkey.

##### Spatial tuning index

We employed two different methods to assess whether the outcome-selective neurons exhibited pronounced spatial clustering.

First, we calculated the Euclidean distances between neurons to assess the presence of clustering among different neuron types within each FOV. Utilizing the suite2p software, we obtained positional coordinates for each neuron within the recording plane. Distances between pairs of neurons were measured based on their positional coordinates within the recording plane, with no regard for their depth. We then examined whether the mean distance between neurons of the same cell type (within-type distance) differed significantly from the mean distance between neurons of different cell types (across-type distance). If the within-type distance was significantly smaller than the across-type distance (P < 0.01, Wilcoxon Ranksun test) in a given FOV, we considered the outcome-selective neurons to be spatially clustered within that 7a subregion.

Second, we calculated a spatial turning index based on both the neural selectivity and spatial position of each neuron. The spatial tuning index is defined as:

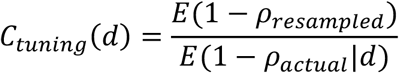

Where *ρ* is the tuning similarity between neuron pairs, and *d* is the spatial distance between neuron pairs. The *ρ* we used equals to the Pearson correlation coefficient of outcome-related activity, calculated using the activity matrix *S* ∈ *R*^*N*×*ct*^ of each neuron. Where *N* denotes the recorded cell number of certain FOV, *c* denotes two outcome conditions (correct and error), and *t* denotes time (1-16 imaging frame after the target offset). The distance *d* was discretized into 50 *μm* bins from 0 to 800 *μm*. The *ρ*_*actual*_ was computed for pairs of neurons whose distances fell within each distance bin. The *ρ*_*resampled*_ was the average tuning similarity between randomly selected cell pairs, calculated using bootstrap (1000 times).

Spatial tuning index values greater than 1 indicate that the outcome selectivity of neurons was more similar than chance level, whereas values smaller than 1 indicate that the outcome selectivity of neurons was more different than chance level. The significance of spatial tuning index is determined by comparing the observed result to a shuffled distribution (bootstrap size = 1000). A result is deemed statistically significant in two circumstances: When the minimum actual C_tuning_ exceeded the values found in 99.9% of the shuffled results, the C_tuning_ values were regarded as significantly higher than chance; when the maximum actual C_tuning_ fell below 99.9% of the shuffled results, the C_tuning_ values were regarded as significantly lower than chance.

##### Demixed principle component analysis (dPCA)

We conducted demixed principal component analysis using the methodology and the code from a previous study^77^ (http://github.com/machenslab/dPCA) to reduce the dimensionality of the population activity as the standard PCA and demixes all task variables. Specifically, we tested how much each task variable (trial outcome, outcome-history, saccade direction, history-saccade interaction, history-outcome interaction, outcome-saccade interaction, timing) contributes to the 7a population activity during the ISAL task.

As demonstrated in the previous study, the dPCA finds separate decoder (F) and encoder (D) matrices for each task variable (∅) by minimizing the loss function:

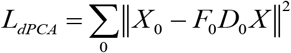

where X is a linear decomposition of the data matrix, which contains the instantaneous activity of the recorded neurons, into variable-specific averages:

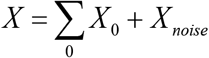

Here, we decomposed the neural activities into seven parts: history dependent (the trial outcome in previous trial), current outcome dependent, saccade-dependent (two saccade directions), dependent on the history-saccade interaction, dependent on the history-outcome interaction, dependent on the outcome-saccade interaction, and noise. The decoder and encoder axes permit us to reduce the data into a few components capturing the majority of the variance of the data dependent on each task variable. Only data from Monkey H were included in this analysis.

#### Quantify the evolution of outcome selectivity during AL

##### Quantify the proportion of neurons that changed selectivity

To evaluate the stability of neural encodings (either outcome encoding or saccade direction encoding) over the AL course, we calculated the proportion of neurons whose selectivity changed from one day to another within the learning course. To achieve this, we first assessed whether each neuron exhibited significant saccade or outcome selectivity. We employed FEV analysis to quantify the strength of neural selectivity across various time windows (using the same time windows used for identifying selective neurons). We determined a neuron’s selectivity preference for a particular day based on the time window with the highest FEV value.

Subsequently, we categorized the neurons in each learning course into several groups. 1) Neurons that exhibited significant selectivity on the familiar day but lost this selectivity across all learning days were classified as ‘lost selectivity’. 2) Neurons that lacked significant selectivity on the familiar day but developed significant selectivity on all learning days were designated as ‘gain selectivity’. 3) Neurons that consistently displayed significant selectivity on every recording day within a learning session without altering their preference were termed ‘persistent selectivity’. 4) Neurons that did not exhibit significant selectivity on any recording day during that learning session were categorized as ‘no selectivity’. 5) The remaining neurons, which encompassed those that altered their outcome selectivity across different learning days, were grouped as ‘change selectivity’.

##### Correlation of neural encoding across different days

In order to quantify the stability of the neural encoding across each learning course, we calculated the correlation of outcome/saccade encoding between two consecutive days within the learning course for each FOV. This process involved two steps. 1) For each individual neuron within the FOV, we quantified the strength of the outcome/saccade encoding using FEV analysis and determined their preferences for each time point. Consequently, for each time point, we constructed an encoding vector that represented the neural encoding within each FOV. This vector incorporated both the strength and preference of neural encodings for all identified neurons within this FOV. 2) At each time point, we calculated the Pearson correlation between the encoding vectors for two consecutive days. This resulted in a correlation value that indicated the similarity of the neural encodings between those two days for each of the given tine point. We repeated the procedures for all time points included in the analysis. In **Figure 6a**, neural encoding correlations were computed separately for two conditions: (1) between the familiar day and the first learning day (familiar-to-novel) and (2) between successive learning days (novel-to-novel). For the novel-to-novel condition, we first calculated pairwise correlations between each pair of successive learning days (while monkeys were acquiring the same novel associations) and then averaged these correlation values.

##### Alignment index

To quantify the relationship between the population codes for trial outcome across different days, we calculated the alignment index (AI), similar as in the previous studies^78^. We quantified the degree of alignment/overlap between the neural subspaces of the population activity from different days within each learning course, and tested how these neural subspaces, responsible for encoding trial outcomes or saccade direction, evolved throughout the learning course.

To calculate AI, we first constructed the population activity matrix *P*_*i*_ ∈ *R*^*N*×*CT*^ for each day, where *N* is the total number of neurons, *C* is the numbers of conditions and *T* is the numbers of time points. Subsequently, to calculate the AI between day *i* and day *j* (*A*(*i, j*)), we conducted principal component analysis (PCA) on activity matrix *P*_*i*_ to obtain the PCs of day *i*, considering each neuron as a variable. We selected the top 10 PCs to form a 10-dimensonal orthogonal axis in N-dimensional neural space. The degree of overlap between neural subspaces can be estimated by projecting the activity matrix from day j onto the PCs of day *i* and quantifying the proportion of variance explained relative to the total variance of population activity on day *j*. Thus, we defined the alignment index *A*(*i, j*) as:

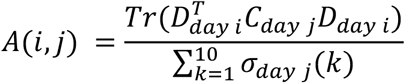

Where *D*_*day i*_ is the top 10 eigenvectors obtained by PCA. *C*_*day j*_ is the covariance of matrix *P*_*j*_. *σ*_*day j*_(*k*) is the *k*-th singular value of *C*_*day j*_. The *Tr*() function denotes the matrix trace, which sums along the diagonal entry. The reverse, *A*(*j, i*), can be calculated in similar ways. The numerator quantifies the proportion of variance in population activity from day j that is captured by the top ten PCs of day i. The denominator normalizes the alignment index with respect to the maximum amount of variance in population activity from day j that can be explained by a ten-dimensional subspace (i.e., the variance captured by the top ten PCs of day j).

As a result, the AI ranged from 0, indicating perfect orthogonality, to 1, signifying perfect alignment.

The detailed parameters used to construct the activity matrix are as follows:

i. To create the activity matrix for saccade encoding (*P*_*saccade*_), we used the activity within a time window spanning from one to16 imaging frames after saccade onset for both saccade directions (left and right).
ii. The activity matrix for outcome encoding (P_outcome_) is constructed using neural activity within the time window spanning from one to 16 imaging frames after target offset for both outcome conditions (correct and error).
iii. The baseline activity matrix (P_baseline_) is assembled using neural activity within the time window spanning from 4 imaging frames before fixation onset to 12 imaging frames after fixation onset, under the same conditions as those used for P_outcome_.
iv. In order to estimate the rate of change in neural encoding within a day/session, we calculate the AIs using data within each day (**Supplementary Fig. 21**). We separated the neural data into two equal-sized portions based on the trial sequence (i.e., the first/second half of all trials, or the odd/even trials). We then constructed the activity matrix for each part of the data.
v. The activity matrix representing changes in outcome or baseline activity across two consecutive days is defined as the difference between the activity matrices for the two days.

In order to quantify the relationship between the outcome encodings on familiar day and learning days, we calculated the *A*(*familar, novel*) using the activity matrix from familiar day and that from each of the learning days for every learning course (**Figure 6b**). The AI values was calculated by projecting the activity matrix from each of the learning days onto the top PCs of the activity matrix from the familiar day. Subsequently, we averaged these AI values to estimate the similarity of neural encodings between familiar day and learning days. Similarly, we calculated the *A*(*novel, novel*) by employing the activity matrix from different learning days for every learning course to quantify the relationship between the outcome encodings on different learning days. The resulting AI values, obtained by projecting the activity matrix from the later learning days onto the top PCs of the activity matrix from the earlier learning days, were averaged to measure the neural encoding similarity between different learning days.

##### Remapping index

We calculated the remapping index as the ratio of inter-day changes in population-level outcome encoding between familiar days and learning days (novel-familiar group) compared to inter-day changes between different learning days (novel-novel group), shown as follows:

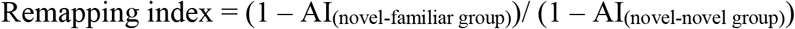

where AI indicates the alignment index quantifying the overlap of the population-level outcome encoding between two days.

#### Quantify the potential mechanisms that constrained the evolution of outcome selectivity during AL

##### Signal correlation

To quantify the similarity of outcome selectivity between different neurons, we calculated the signal correlation for each pair of neurons. In contrast to classical methods, we incorporated time with outcome conditions to calculate the signal correlation. To achieve this, we first constructed the activity vector A∈ *v*^*ct*^, where c denotes two outcome conditions (correct and error), and t denotes time (1-16 imaging frame after the target offset). The signal correlation between each pair of neurons was defined as the Pearson correlation between their respective activity vectors.

##### Noise correlation

We calculated the noise correlation between different neurons to quantify the connectivity between them following these steps: 1) Single-trial responses were estimated using the averaged activity within two time windows (1-4 frames or 5-8 frames after target offset) in the ISAL task. Alternatively, another time window (1-4 frames or 5-8 frames before cue onset) was used to calculate noise correlation, which yielded quantitatively similar results.2) Pearson correlations were calculated separately for correct trials and error trials using the pairwise trial-by-trial activity. 3) We calculated a single correlation value by averaging the two correlation values for correct and error trials, using the trial numbers of each condition as weights.

For each data session, we obtained two correlation matrixes *C*_*noise*_, *C*_*sittnal*_ ∈ *R*^*N*×*N*^, within N representing the number of neurons within each FOV. In **Figure 7b**, in order to quantify the correlation between noise correlation and signal correlation, we transformed the upper-triangles of those matrixes into 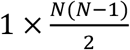 vectors, and subsequently calculated the correlations between these vectors from different days. In **Figure 7c**, we defined changes in correlations between two days as the difference between the vectors from those different days.

In **Figure 7d-g**, we calculated the within-type correlations in the current day, and tested whether these noise correlations could predict changes in outcome selectivity on the following day. Specifically, this analysis focused on neurons belonging to the same type (CN, EN, or non-selective) on the current day but transitioning to different types on the next day. We categorized these neurons into ‘within-future-type’ pairs (neurons belonging to the same type on the next day) and ‘cross-future-type’ pairs (neurons transitioning to the different types on the next day) based on their outcome-selectivity on the next day (**Figure 7d**). In **Figure 7h-k**, we tested whether the noise correlation among cross-type neuron pairs could also predict changes in outcome selectivity during AL. Here, we focused on the neurons that were initially belonged to different types on the current day but converged to the same type on the following day. Similar to the previous analysis, we categorized these neurons into ‘within-future-type’ pairs and ‘cross-future-type’ pairs, based on their outcome selectivity on the next day (**Figure 7i)**. For both analyses, only the data sessions in which there were sufficient number of neurons (n>=10) in each neuron type were included. Due to limited data sessions meeting this neuron number requirement for all cell types, we independently selected data sessions for outcome-selective and non-selective neurons for the analysis.

In **Figure 7g-k**, we also employed the signal correlation to predict changes in outcome selectivity during AL. The calculation steps were identical to those used for noise correlation, with the distinction that we utilized signal correlation between neurons to predict changes in neural ensembles.

## Notes

### Competing Interest Statement

The authors have declared no competing interest.

## References

1 Ullsperger, M., Danielmeier, C. & Jocham, G. Neurophysiology of performance monitoring and adaptive behavior. Physiol Rev 94, 35–79, doi:10.1152/physrev.00041.2012 (2014).

2 Lee, D., Seo, H. & Jung, M. W. Neural basis of reinforcement learning and decision making. Annu Rev Neurosci 35, 287–308, doi:10.1146/annurev-neuro-062111-150512 (2012).

3 Histed, M. H., Pasupathy, A. & Miller, E. K. Learning substrates in the primate prefrontal cortex and striatum: sustained activity related to successful actions. Neuron 63, 244–253, doi:10.1016/j.neuron.2009.06.019 (2009).

4 Heilbronner, S. R. & Platt, M. L. Causal evidence of performance monitoring by neurons in posterior cingulate cortex during learning. Neuron 80, 1384–1391, doi:10.1016/j.neuron.2013.09.028 (2013).

5 Sarafyazd, M. & Jazayeri, M. Hierarchical reasoning by neural circuits in the frontal cortex. Science 364, doi:10.1126/science.aav8911 (2019).

6 Sendhilnathan, N., Semework, M., Goldberg, M. E. & Ipata, A. E. Neural Correlates of Reinforcement Learning in Mid-lateral Cerebellum. Neuron 106, 188–198 e185, doi:10.1016/j.neuron.2019.12.032 (2020).

7 Seo, H., Barraclough, D. J. & Lee, D. Dynamic signals related to choices and outcomes in the dorsolateral prefrontal cortex. Cereb Cortex 17 Suppl 1, i110–117, doi:10.1093/cercor/bhm064 (2007).

8 Wirth, S. et al. Trial outcome and associative learning signals in the monkey hippocampus. Neuron 61, 930–940, doi:10.1016/j.neuron.2009.01.012 (2009).

9 Barraclough, D. J., Conroy, M. L. & Lee, D. Prefrontal cortex and decision making in a mixed-strategy game. Nat Neurosci 7, 404–410, doi:10.1038/nn1209 (2004).

10 Seo, H., Cai, X., Donahue, C. H. & Lee, D. Neural correlates of strategic reasoning during competitive games. Science 346, 340–343, doi:10.1126/science.1256254 (2014).

11 Asaad, W. F., Rainer, G. & Miller, E. K. Neural activity in the primate prefrontal cortex during associative learning. Neuron 21, 1399–1407, doi:10.1016/s0896-6273(00)80658-3 (1998).

12 Wirth, S. et al. Single neurons in the monkey hippocampus and learning of new associations. Science 300, 1578–1581, doi:10.1126/science.1084324 (2003).

13 Chen, L. L. & Wise, S. P. Supplementary eye field contrasted with the frontal eye field during acquisition of conditional oculomotor associations. J Neurophysiol 73, 1122–1134, doi:10.1152/jn.1995.73.3.1122 (1995).

14 Mitz, A. R., Godschalk, M. & Wise, S. P. Learning-dependent neuronal activity in the premotor cortex: activity during the acquisition of conditional motor associations. J Neurosci 11, 1855–1872, doi:10.1523/JNEUROSCI.11-06-01855.1991 (1991).

15 Pasupathy, A. & Miller, E. K. Different time courses of learning-related activity in the prefrontal cortex and striatum. Nature 433, 873–876, doi:10.1038/nature03287 (2005).

16 Messinger, A., Squire, L. R., Zola, S. M. & Albright, T. D. Neuronal representations of stimulus associations develop in the temporal lobe during learning. Proceedings of the National Academy of Sciences of the United States of America 98, 12239–12244, doi:10.1073/pnas.211431098 (2001).

17 Brasted, P. J. & Wise, S. P. Comparison of learning-related neuronal activity in the dorsal premotor cortex and striatum. The European journal of neuroscience 19, 721–740, doi:10.1111/j.0953-816x.2003.03181.x (2004).

18 Lee, D. & Seo, H. Mechanisms of reinforcement learning and decision making in the primate dorsolateral prefrontal cortex. Ann N Y Acad Sci 1104, 108–122, doi:10.1196/annals.1390.007 (2007).

19 Cox, J. & Witten, I. B. Striatal circuits for reward learning and decision-making. Nature reviews. Neuroscience 20, 482–494, doi:10.1038/s41583-019-0189-2 (2019).

20 Averbeck, B. & O’Doherty, J. P. Reinforcement-learning in fronto-striatal circuits. Neuropsychopharmacology 47, 147–162, doi:10.1038/s41386-021-01108-0 (2022).

21 Rushworth, M. F., Noonan, M. P., Boorman, E. D., Walton, M. E. & Behrens, T. E. Frontal cortex and reward-guided learning and decision-making. Neuron 70, 1054–1069, doi:10.1016/j.neuron.2011.05.014 (2011).

22 Packard, M. G. & Knowlton, B. J. Learning and memory functions of the Basal Ganglia. Annu Rev Neurosci 25, 563–593, doi:10.1146/annurev.neuro.25.112701.142937 (2002).

23 Makino, H., Hwang, E. J., Hedrick, N. G. & Komiyama, T. Circuit Mechanisms of Sensorimotor Learning. Neuron 92, 705–721, doi:10.1016/j.neuron.2016.10.029 (2016).

24 Murray, E. A. & Rudebeck, P. H. Specializations for reward-guided decision-making in the primate ventral prefrontal cortex. Nature reviews. Neuroscience 19, 404–417, doi:10.1038/s41583-018-0013-4 (2018).

25 Ni, A. M., Ruff, D. A., Alberts, J. J., Symmonds, J. & Cohen, M. R. Learning and attention reveal a general relationship between population activity and behavior. Science 359, 463–465, doi:10.1126/science.aao0284 (2018).

26 Freedman, D. J. & Assad, J. A. Experience-dependent representation of visual categories in parietal cortex. Nature 443, 85–88, doi:10.1038/nature05078 (2006).

27 Law, C. T. & Gold, J. I. Neural correlates of perceptual learning in a sensory-motor, but not a sensory, cortical area. Nat Neurosci 11, 505–513, doi:10.1038/nn2070 (2008).

28 Morrison, S. E., Saez, A., Lau, B. & Salzman, C. D. Different time courses for learning-related changes in amygdala and orbitofrontal cortex. Neuron 71, 1127–1140, doi:10.1016/j.neuron.2011.07.016 (2011).

29 Gold, J. I. & Shadlen, M. N. The neural basis of decision making. Annu Rev Neurosci 30, 535–574, doi:10.1146/annurev.neuro.29.051605.113038 (2007).

30 Huk, A. C., Katz, L. N. & Yates, J. L. The Role of the Lateral Intraparietal Area in (the Study of) Decision Making. Annu Rev Neurosci 40, 349–372, doi:10.1146/annurev-neuro-072116-031508 (2017).

31 Platt, M. L. & Glimcher, P. W. Neural correlates of decision variables in parietal cortex. Nature 400, 233–238, doi:10.1038/22268 (1999).

32 Bisley, J. W. & Goldberg, M. E. Attention, intention, and priority in the parietal lobe. Annu Rev Neurosci 33, 1–21, doi:10.1146/annurev-neuro-060909-152823 (2010).

33 Colby, C. L. & Goldberg, M. E. Space and attention in parietal cortex. Annu Rev Neurosci 22, 319–349, doi:10.1146/annurev.neuro.22.1.319 (1999).

34 Andersen, R. A. & Buneo, C. A. Intentional maps in posterior parietal cortex. Annu Rev Neurosci 25, 189–220, doi:10.1146/annurev.neuro.25.112701.142922 (2002).

35 Zhou, Y. & Freedman, D. J. Posterior parietal cortex plays a causal role in perceptual and categorical decisions. Science 365, 180–185, doi:10.1126/science.aaw8347 (2019).

36 Zhou, Y., Liu, Y. & Zhang, M. Neuronal Correlates of Many-To-One Sensorimotor Mapping in Lateral Intraparietal Cortex. Cereb Cortex, doi:10.1093/cercor/bhaa145 (2020).

37 Seo, H., Barraclough, D. J. & Lee, D. Lateral intraparietal cortex and reinforcement learning during a mixed-strategy game. J Neurosci 29, 7278–7289, doi:10.1523/JNEUROSCI.1479-09.2009 (2009).

38 Felleman, D. J. & Van Essen, D. C. Distributed hierarchical processing in the primate cerebral cortex. Cereb Cortex 1, 1–47, doi:10.1093/cercor/1.1.1-a (1991).

39 Andersen, R. A., Snyder, L. H., Bradley, D. C. & Xing, J. Multimodal representation of space in the posterior parietal cortex and its use in planning movements. Annu Rev Neurosci 20, 303–330, doi:10.1146/annurev.neuro.20.1.303 (1997).

40 Gnadt, J. W. & Andersen, R. A. Memory related motor planning activity in posterior parietal cortex of macaque. Exp Brain Res 70, 216–220 (1988).

41 Constantinidis, C. & Steinmetz, M. A. Posterior parietal cortex automatically encodes the location of salient stimuli. J Neurosci 25, 233–238, doi:10.1523/JNEUROSCI.3379-04.2005 (2005).

42 Andersen, R. A., Essick, G. K. & Siegel, R. M. Encoding of spatial location by posterior parietal neurons. Science 230, 456–458, doi:10.1126/science.4048942 (1985).

43 Snyder, L. H., Grieve, K. L., Brotchie, P. & Andersen, R. A. Separate body- and world-referenced representations of visual space in parietal cortex. Nature 394, 887–891, doi:10.1038/29777 (1998).

44 Chafee, M. V., Crowe, D. A., Averbeck, B. B. & Georgopoulos, A. P. Neural correlates of spatial judgement during object construction in parietal cortex. Cereb Cortex 15, 1393–1413, doi:10.1093/cercor/bhi021 (2005).

45 Lewis, J. W. & Van Essen, D. C. Corticocortical connections of visual, sensorimotor, and multimodal processing areas in the parietal lobe of the macaque monkey. J Comp Neurol 428, 112–137, doi:10.1002/1096-9861(20001204)428:1<112::aid-cne8>3.0.co;2-9 (2000).

46 Rockland, K. S. & Van Hoesen, G. W. Some temporal and parietal cortical connections converge in CA1 of the primate hippocampus. Cerebral Cortex 9, 232–237, doi:DOI 10.1093/cercor/9.3.232 (1999).

47 Cavada, C. & Goldman-Rakic, P. S. Posterior parietal cortex in rhesus monkey: II. Evidence for segregated corticocortical networks linking sensory and limbic areas with the frontal lobe. J Comp Neurol 287, 422–445, doi:10.1002/cne.902870403 (1989).

48 Roy, N. A. et al. Extracting the dynamics of behavior in sensory decision-making experiments. Neuron 109, 597–610 e596, doi:10.1016/j.neuron.2020.12.004 (2021).

49 Alonso, J. M. & Martinez, L. M. Functional connectivity between simple cells and complex cells in cat striate cortex. Nature Neuroscience 1, 395–403, doi:Doi 10.1038/1609 (1998).

50 Cohen, M. R. & Kohn, A. Measuring and interpreting neuronal correlations. Nat Neurosci 14, 811–819, doi:10.1038/nn.2842 (2011).

51 Daw, N. D., O’Doherty, J. P., Dayan, P., Seymour, B. & Dolan, R. J. Cortical substrates for exploratory decisions in humans. Nature 441, 876–879, doi:10.1038/nature04766 (2006).

52 Donoso, M., Collins, A. G. & Koechlin, E. Human cognition. Foundations of human reasoning in the prefrontal cortex. Science 344, 1481–1486, doi:10.1126/science.1252254 (2014).

53 Koob, G. F., Stinus, L., Le Moal, M. & Bloom, F. E. Opponent process theory of motivation: neurobiological evidence from studies of opiate dependence. Neurosci Biobehav Rev 13, 135–140, doi:10.1016/s0149-7634(89)80022-3 (1989).

54 Petrides, M. Visuo-motor conditional associative learning after frontal and temporal lesions in the human brain. Neuropsychologia 35, 989–997, doi:10.1016/s0028-3932(97)00026-2 (1997).

55 Rothenhoefer, K. M. et al. Effects of Ventral Striatum Lesions on Stimulus-Based versus Action-Based Reinforcement Learning. J Neurosci 37, 6902–6914, doi:10.1523/JNEUROSCI.0631-17.2017 (2017).

56 Rudebeck, P. H., Saunders, R. C., Prescott, A. T., Chau, L. S. & Murray, E. A. Prefrontal mechanisms of behavioral flexibility, emotion regulation and value updating. Nat Neurosci 16, 1140–1145, doi:10.1038/nn.3440 (2013).

57 Taswell, C. A., Costa, V. D., Murray, E. A. & Averbeck, B. B. Ventral striatum’s role in learning from gains and losses. Proceedings of the National Academy of Sciences of the United States of America 115, E12398–E12406, doi:10.1073/pnas.1809833115 (2018).

58 Akrami, A., Kopec, C. D., Diamond, M. E. & Brody, C. D. Posterior parietal cortex represents sensory history and mediates its effects on behaviour. Nature 554, 368-+, doi:10.1038/nature25510 (2018).

59 Hwang, E. J., Dahlen, J. E., Mukundan, M. & Komiyama, T. History-based action selection bias in posterior parietal cortex. Nat Commun 8, 1242, doi:10.1038/s41467-017-01356-z (2017).

60 Glascher, J., Daw, N., Dayan, P. & O’Doherty, J. P. States versus rewards: dissociable neural prediction error signals underlying model-based and model-free reinforcement learning. Neuron 66, 585–595, doi:10.1016/j.neuron.2010.04.016 (2010).

61 Xue, C., Kramer, L. E. & Cohen, M. R. Dynamic task-belief is an integral part of decision-making. Neuron 110, 2503–2511 e2503, doi:10.1016/j.neuron.2022.05.010 (2022).

62 Andersen, R. A., Essick, G. K. & Siegel, R. M. Neurons of area 7 activated by both visual stimuli and oculomotor behavior. Exp Brain Res 67, 316–322, doi:10.1007/BF00248552 (1987).

63 Barash, S., Bracewell, R. M., Fogassi, L., Gnadt, J. W. & Andersen, R. A. Saccade-related activity in the lateral intraparietal area. I. Temporal properties; comparison with area 7a. J Neurophysiol 66, 1095–1108, doi:10.1152/jn.1991.66.3.1095 (1991).

64 Andersen, R. A., Bracewell, R. M., Barash, S., Gnadt, J. W. & Fogassi, L. Eye position effects on visual, memory, and saccade-related activity in areas LIP and 7a of macaque. J Neurosci 10, 1176–1196, doi:10.1523/JNEUROSCI.10-04-01176.1990 (1990).

65 Snyder, L. H., Batista, A. P. & Andersen, R. A. Coding of intention in the posterior parietal cortex. Nature 386, 167–170, doi:10.1038/386167a0 (1997).

66 Heider, B. & Siegel, R. M. Optical imaging of visually guided reaching in macaque posterior parietal cortex. Brain Struct Funct 219, 495–509, doi:10.1007/s00429-013-0513-y (2014).

67 Fyhn, M., Hafting, T., Treves, A., Moser, M. B. & Moser, E. I. Hippocampal remapping and grid realignment in entorhinal cortex. Nature 446, 190–194, doi:10.1038/nature05601 (2007).

68 Colgin, L. L., Moser, E. I. & Moser, M. B. Understanding memory through hippocampal remapping. Trends in Neurosciences 31, 469–477, doi:10.1016/j.tins.2008.06.008 (2008).

69 Azeredo da Silveira, R. & Rieke, F. The Geometry of Information Coding in Correlated Neural Populations. Annu Rev Neurosci 44, 403–424, doi:10.1146/annurev-neuro-120320-082744 (2021).

70 Jeanne, J. M., Sharpee, T. O. & Gentner, T. Q. Associative learning enhances population coding by inverting interneuronal correlation patterns. Neuron 78, 352–363, doi:10.1016/j.neuron.2013.02.023 (2013).

71 Averbeck, B. B., Latham, P. E. & Pouget, A. Neural correlations, population coding and computation. Nature Reviews Neuroscience 7, 358–366, doi:10.1038/nrn1888 (2006).

72 Li, M., Liu, F., Jiang, H., Lee, T. S. & Tang, S. Long-Term Two-Photon Imaging in Awake Macaque Monkey. Neuron 93, 1049–1057 e1043, doi:10.1016/j.neuron.2017.01.027 (2017).

73 Tang, S. et al. Complex Pattern Selectivity in Macaque Primary Visual Cortex Revealed by Large-Scale Two-Photon Imaging. Curr Biol 28, 38–48 e33, doi:10.1016/j.cub.2017.11.039 (2018).

74 Asaad, W. F., Santhanam, N., McClellan, S. & Freedman, D. J. High-performance execution of psychophysical tasks with complex visual stimuli in MATLAB. J Neurophysiol 109, 249–260, doi:10.1152/jn.00527.2012 (2013).

75 Zhou, Y., Mohan, K. & Freedman, D. J. Abstract Encoding of Categorical Decisions in Medial Superior Temporal and Lateral Intraparietal Cortices. J Neurosci 42, 9069–9081, doi:10.1523/JNEUROSCI.0017-22.2022 (2022).

76 Zhou, Y. et al. Distributed functions of prefrontal and parietal cortices during sequential categorical decisions. Elife 10, doi:10.7554/eLife.58782 (2021).

77 Kobak, D. et al. Demixed principal component analysis of neural population data. Elife 5, doi:10.7554/eLife.10989 (2016).

78 Elsayed, G. F., Lara, A. H., Kaufman, M. T., Churchland, M. M. & Cunningham, J. P. Reorganization between preparatory and movement population responses in motor cortex. Nature Communications 7, doi:ARTN 13239 10.1038/ncomms13239 (2016).

