## Supplementary Fig. 1-21 and Supplementary Table 1-2 for "Neural Correlates of Trial Outcome Monitoring during Long-term Learning in Primate Posterior Parietal Cortex"

**The file includes:**

Supplementary Fig. 1-21
Supplementary Table 1-2

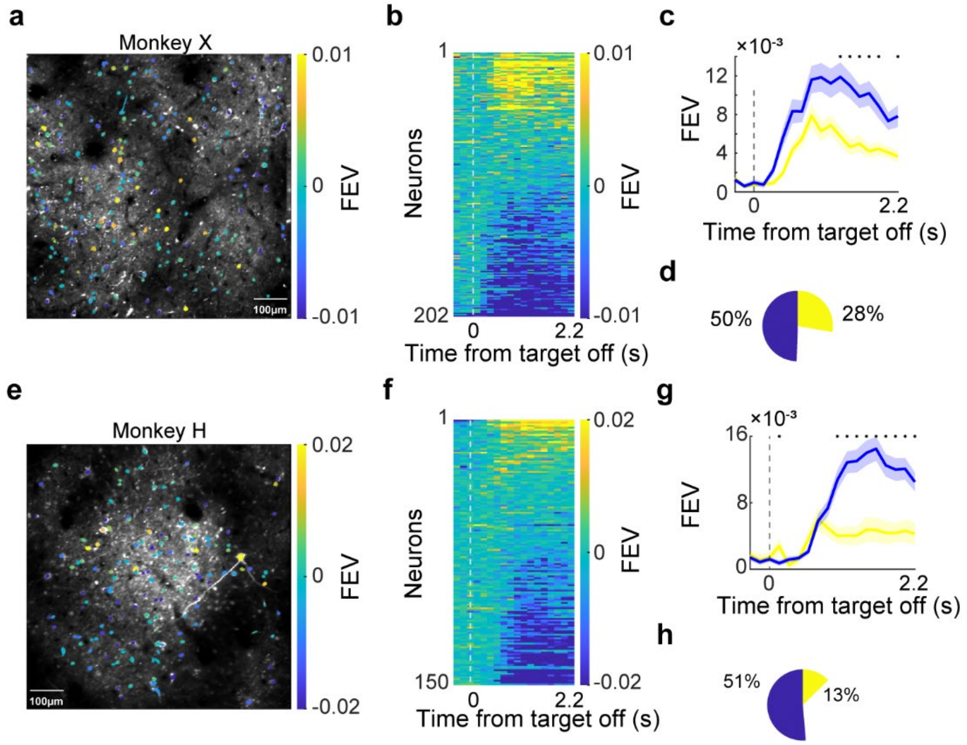


**Supplementary Fig. 1 7a neurons encoded trial outcome during AL. a-d**. An example FOV within which neurons exhibited pronounced outcome selectivity. **a**. An example FOV with identified ROIs. The color bar marks the averaged strength (FEV value) and preference of outcome selectivity for each identified neuron. **b**. The outcome-selectivity for all identified neurons in the example FOV was shown as a function of time. **c**. The averaged magnitude of outcome selectivity was shown separately for both CNs and ENs. Shaded region denotes SEM. The stars denote the time points with significant difference between CNs and ENs ($P < 0.01$, Wilcoxon test). **d**. The percentages of CNs and ENs among all identified neurons within this FOV. **e-h**. Another example FOV which was shown in the same format as in **a-d**.

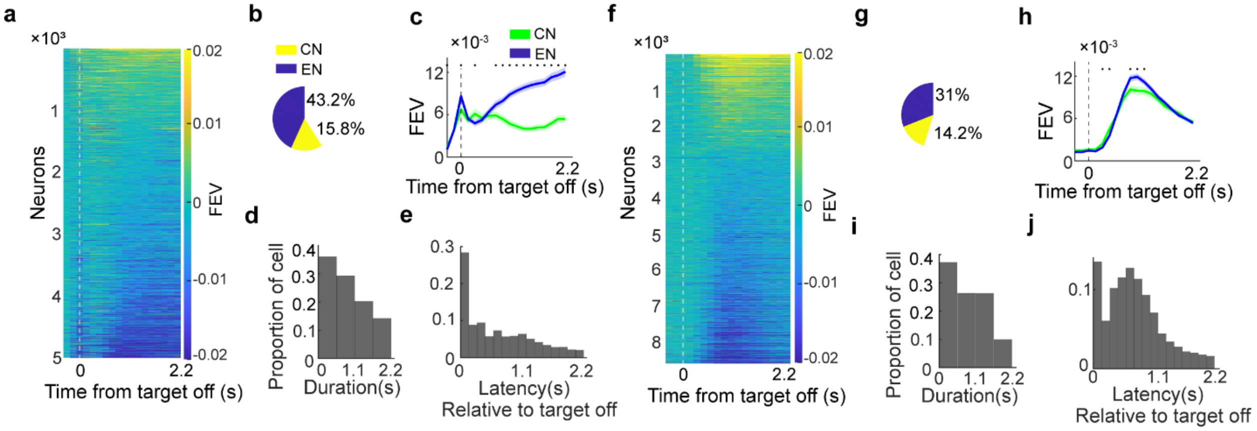


**Supplementary Fig. 2 Outcome encoding of 7a neurons in Monkey H and Monkey X. a**. Outcome selectivity for identified outcome-selective neurons in monkey H. **b**. The percentages of CNs and ENs among all identified neurons. **c**. The averaged magnitude of outcome selectivity were shown separately for both CNs and ENs. Shaded region denotes SEM. **d**. Duration of outcome encoding for outcome-selective neurons. Each time window spans 0.54s. **e**. Latency of outcome selectivity relative to target offset for the outcome-selective neurons. **f-j**. Outcome selectivity for identified outcome-selective neurons in monkey X, shown in the same format as **a -e**.

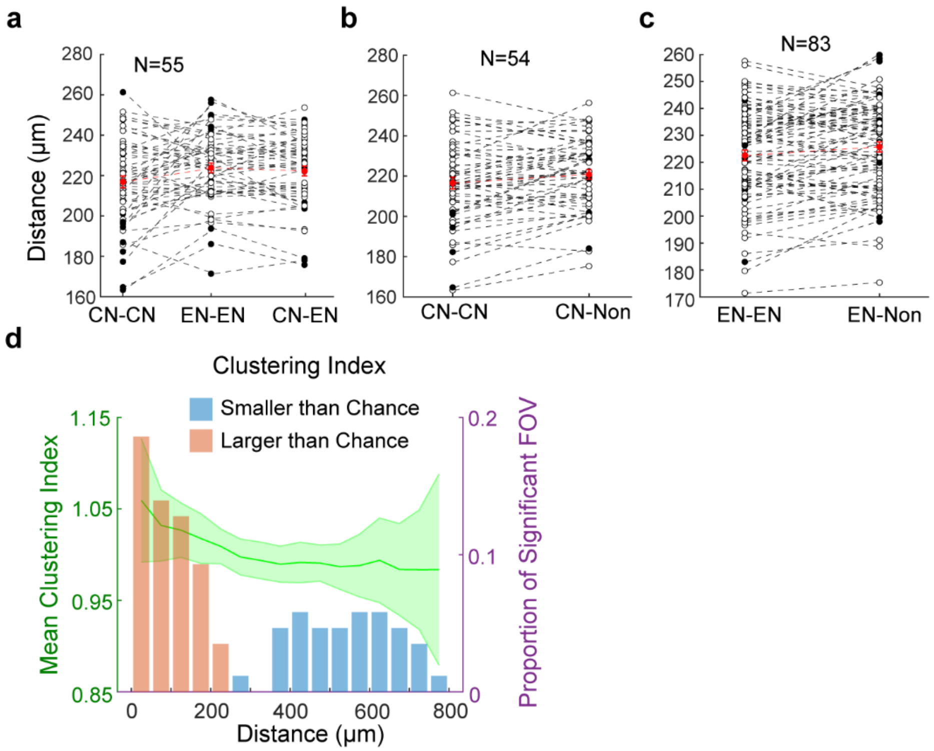


**Supplementary Fig. 3 The outcome selective neurons did not show obvious spatial clustering in 7a. a-c**. Averaged distances between different pairs of neurons shown separately for different neuron types. Distances between pairs of neurons were measured based on their positional coordinates within the recording plane, with no regard for their depth. **a**. Comparisons of distance between different types of outcome selective neurons. Neuron pairs within the same neuron type (CN-CN and EN-EN) and neuron pairs of different neuron types were shown separately. Dashed lines connect data within each recording session. Open circles represent the data sessions where there was no significant difference in the comparison, while filled dots denote the data session there was significant difference (Wilcoxon test, $P < 0.01$). Red bars denote the mean with SEM for each group. **b.** Comparisons of the distances between neuron pairs from CNs and the distance between CNs and non-selective neurons. **c.** Comparisons of distances between neuron pairs from ENs and the distance between ENs and non-selective neurons. Spatial clustering of outcome-selective neurons was observed in very few sessions. **d.** The spatial tuning index for all recording sessions. We calculated the spatial turning index based on both neural selectivity and spatial position of each identified neuron. Spatial tuning index values greater than 1 indicate that the outcome selectivity of neurons was more similar than chance level, whereas values smaller than 1 indicate that the outcome selectivity of neurons was more different than chance level. The spatial tuning index was computed for pairs of neurons whose distances fell within each distance bin (50 μm bins from 0 to 800 μm). The averaged mean and SEM of spatial tuning index values (green) across all recording sessions were overlaid with the proportions of sessions that exhibited significant spatial clustering (red and blue bars). The left Y-axis represents spatial tuning index values, while the right Y-axis represents the proportion of sessions where the spatial tuning index significantly differed from chance level. Only a small proportion of data sessions exhibited significant spatial clustering for outcome encoding.

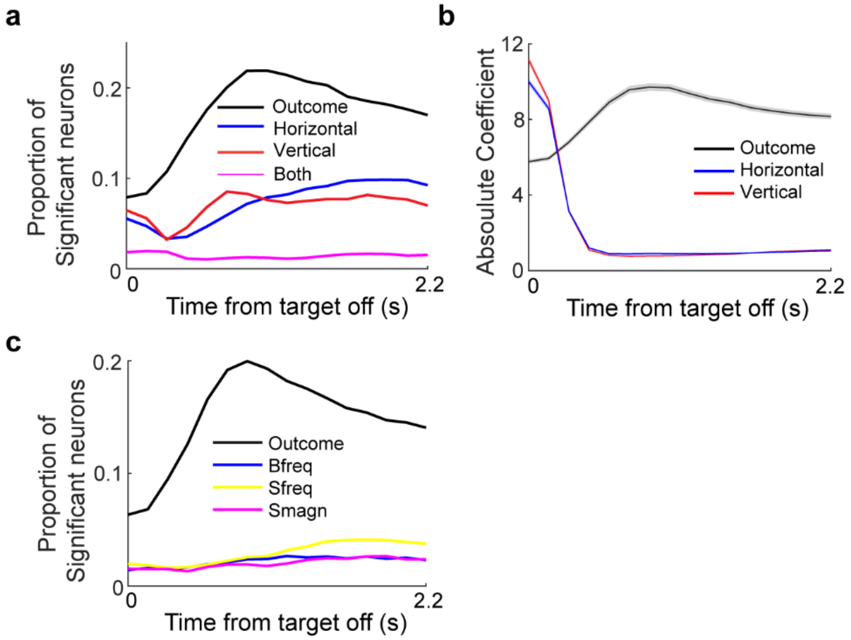


**Supplementary Fig. 4 The outcome selectivity in 7a cannot be simply attributed to differences in eye position and eye movements between correct and error trials.** To assess whether the outcome selectivity of 7a neurons primarily stems from encodings relative to monkeys' eye positions during the inter-trial interval, we employed a linear regression model to quantify the dependence of 7a neuronal activity on monkeys' eye positions in both horizontal and vertical directions, as well as trial outcome. **a**. The proportion of neurons that exhibited a significant correlation between monkeys' eye positions and their activity. The linear regression was applied to neuronal activity for each time frame after target offset. **b**. The resulted absolute coefficient values from the linear regression for all 7a neurons. The absolute coefficient values for monkeys' eye positions in both directions rapidly diminished shortly after the saccade target offset. In contrast, the absolute coefficient values for trial outcomes exhibited the opposite trend. Shaded region denotes SEM. The monkeys' eye positions during inter-trial interval did not contribute primarily to the neuronal activity in area 7a. Furthermore, to determine whether the outcome selectivity of 7a neurons was primarily contributed by differences in the monkeys' saccadic eye movements and blinks during the inter-trial interval, we employed an additional linear regression model. This model quantified the relationship between 7a neuronal activity and variables such as saccade frequency, saccade amplitude, eye blink frequency, and trial outcome. **c**. The proportion of neurons that exhibited significant encoding of trial outcome, eye blink frequency, saccade frequency, and saccade amplitude for each frame after target offset. (Bfreq: blink frequency, Sfreq: saccade frequency, Smagn: saccade magnitude).

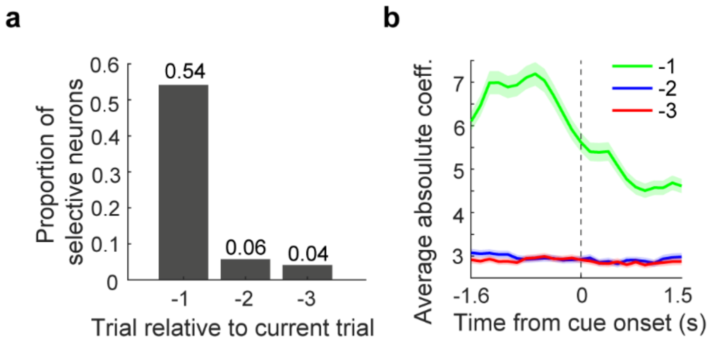


**Supplementary Fig. 5 7a neurons encoded trial outcome in the previous trials. a.** The proportions of neurons that significantly encoded the trial outcome in the trial occurring one, two, or three trials before the current trial. **b.** A linear regression model was employed to quantify the strength of the encoding of outcome history, and the absolute coefficients of the model fitting are presented for the encoding of outcomes in trials occurring one, two, or three trials before the current trial. Shaded region denotes SEM.

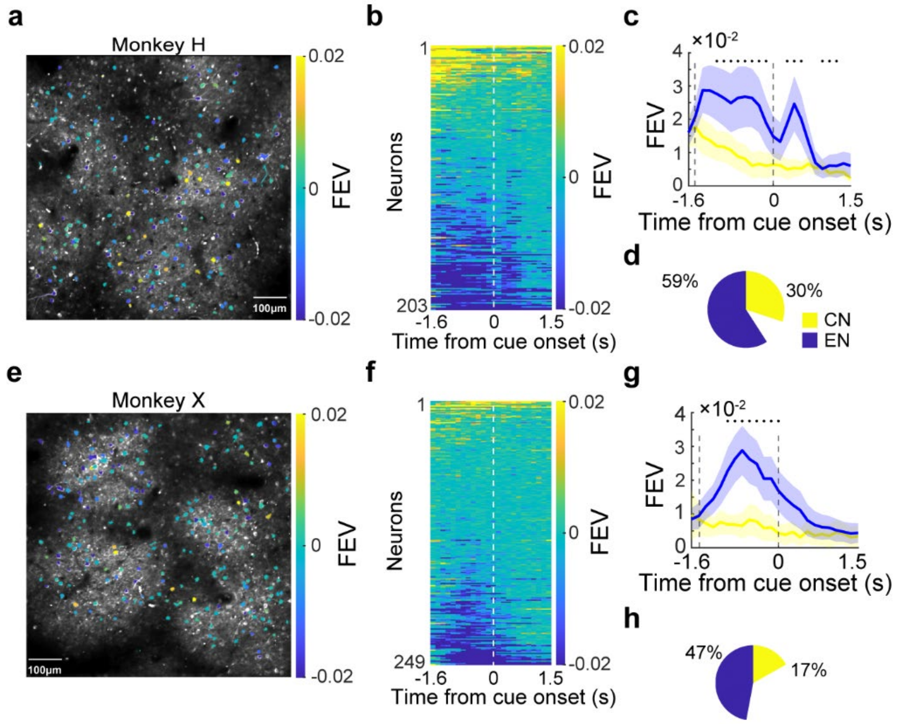


**Supplementary Fig. 6 Encoding of trial outcome in the previous trial. a-d**. An example FOV within which neurons exhibited pronounced encoding of the trial outcome in the previous trial (outcome history). **a.** An example FOV with identified ROIs. The color bar marks the averaged strength (FEV value) and preference of outcome-history encoding for each identified neuron. **b.** The outcome-history selectivity for all identified neurons in the example FOV was shown as a function of time. **c.** The averaged magnitude of outcome-history selectivity were shown separately for CNs and ENs. Shaded region denotes SEM. The stars denote the time points with significant difference between CNs and ENs ($P < 0.01$, Wilcoxon test). **d.** The percentages of CNs and ENs among all identified neurons within this FOV. **e-h**. Another example FOV presented in the same format as in a-d.

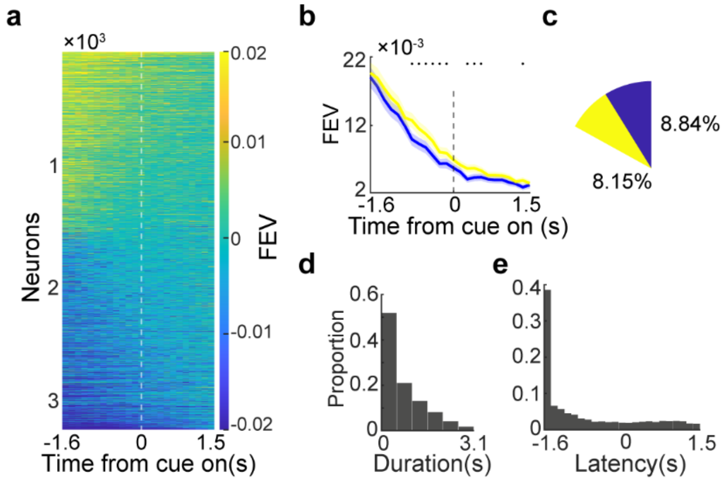


**Supplementary Fig 7. Outcome-history encoding of 7a neurons in Monkey X.** Monkey X exhibited a high frequency of fix-break trials during task performance (0.5660 ± 0.080), resulting in an excessive number of fix-break trials interspersed between completed trials. Consequently, we excluded this monkey's data from the analysis of outcome history encoding in **Figure 2n-r**. **a.** The outcome-history selectivity of all identified neurons that showed significant encoding on outcome history (3229 of 19008) across all FOVs from this monkey was displayed over time. **b.** The averaged magnitude of outcome selectivity were shown separately for both CNs and ENs. Shaded region denotes SEM. **c.** The percentages of CNs and ENs among all identified neurons. **d.** The duration of outcome-history encoding for all outcome-history selective neurons. **e.** The latency of outcome selectivity relative to fixation onset for all the PO selective neurons.

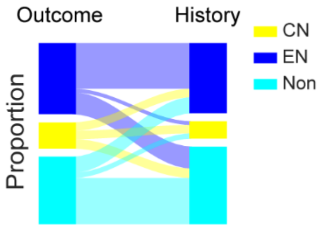


**Supplementary Fig. 8 The PO encoding was not merely a continuation of the outcome encoding from the previous trial.** A significant fraction of neurons exhibited inconsistent encodings of trial outcome or outcome history.

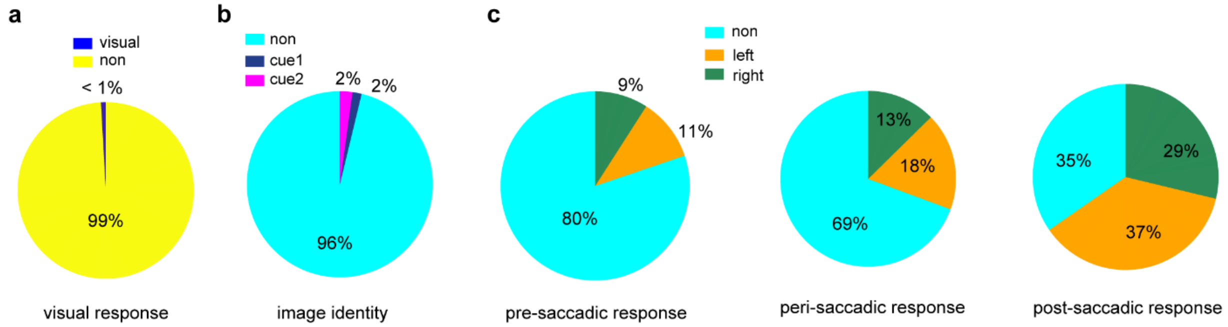


**Supplementary Fig 9. Neuronal encoding of 7a neurons in different task epochs. a.** The percentages of neurons which showed obvious visual response to the visual cue during the ISAL task. **b.** The percentages of neurons which encoded cue selectivity during cue presentation interval of the ISAL task. **c.** The percentages of neurons which encoded saccade direction during pre-saccadic interval (delay period), peri saccadic interval and post saccadic interval of the ISAL task.

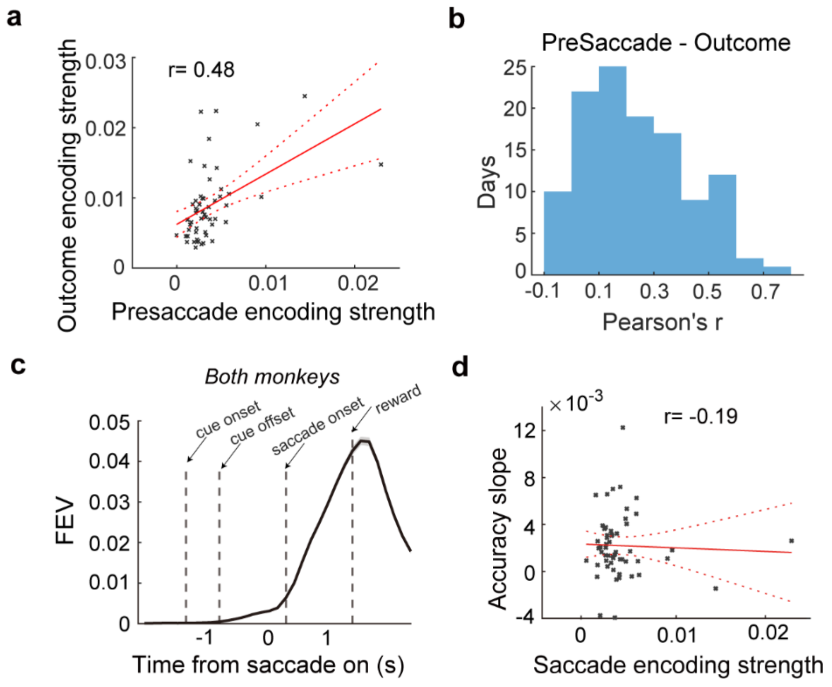


**Supplementary Fig. 10 Saccade direction encoding during the delay period. a.** Correlation between pre-saccade encoding strength and outcome encoding strength. Each dot denotes the averaged results for one FOV. The red line indicates the linear regression fit; dashed lines denote the 95% confidence interval. **b.** The distribution of Correlation values between outcome encoding and saccade direction encoding across all recording sessions. **c.** Time course of saccade direction encoding strength (quantified by FEV) for all saccade-selective neurons. Shaded regions denote SEMs. **d.** Correlation between the averaged strength of saccade direction encoding and the monkeys' performance accuracy during learning.

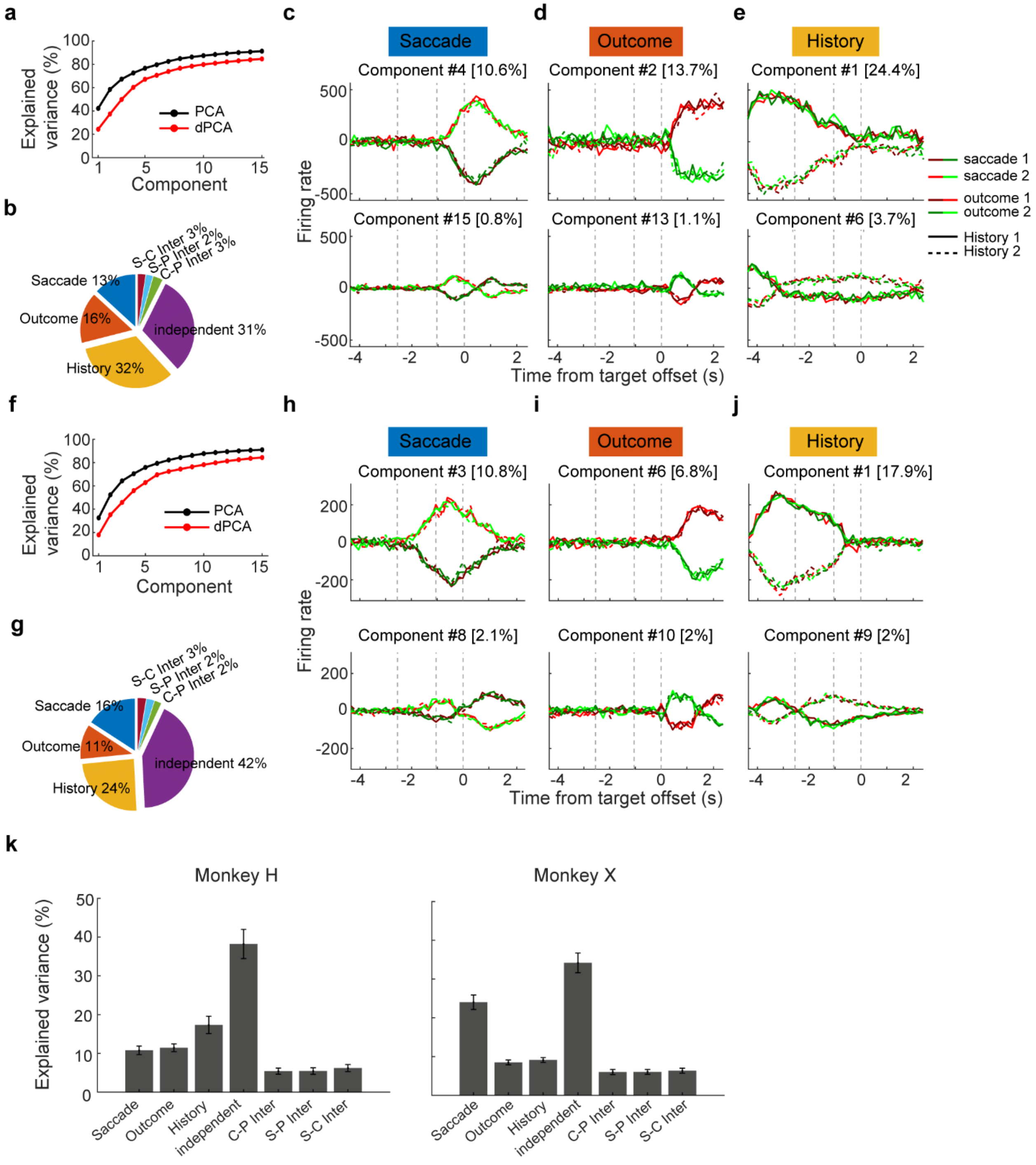


**Supplementary Fig. 11 The strength of encodings of different task variables in 7a were quantified by a demixed principle component analysis (dPCA). a-e.** The dPCA results for one example session. **a.** The accumulated variance explained by top 15 principle components (PCs) in normal PCA analysis, as well as accumulated variance explained by top 15 demixed PCs. **b.** The total variance explained by each task variable for 7a population activity in this FOV. **c-e.** Demixed principal components. The first row: the first demixed PCs of 7a population activity corresponding to saccade direction, trial outcome and outcome history. The second row: the second demixed PCs of 7a population activity. In each subplot, LIP population activity is projected onto the respective dPCA decoder axis, so that there are 8 lines corresponding to 8 conditions (2 trial outcomes × 2 outcome history × 2 saccade directions). Different colors represent different outcome conditions. Different shades represent different saccade directions. Solid and dashed lines represent different outcome-history conditions. The three vertical dashed line mark the time of cue onset, saccade onset, and target offset. **f-j.** The dPCA results for another example session. **k.** The average variance explained by each task variable across all tested FOVs. The error bar denotes SEM. Data from the two monkeys are shown separately in the left and right panels.

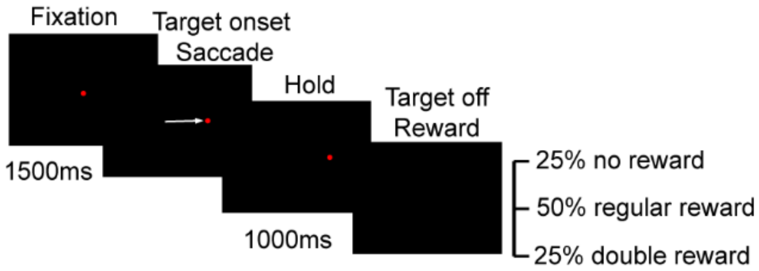


**Supplementary Fig. 12 Schematic illustration of the visual saccade task.** This task was designed as a control task to investigate whether the outcome encoding in PPC solely reflected reward reception. After the monkey acquired central fixation for 1500 ms, the fixation point disappeared, and a visual target appeared at one of four possible locations. Monkeys were required to execute a single saccade to the target within 400 ms and maintain fixation on the target for 1000 ms to receive a juice reward. In one-quarter of the randomly selected trials, monkeys received a double-sized reward, while in another quarter of the randomly selected trials, the juice reward was omitted.

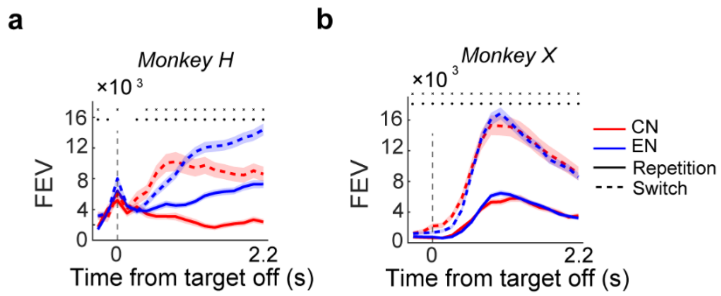


**Supplementary Fig. 13 Outcome history influenced outcome encoding. a-b**. Comparison of average outcome selectivity between trials following outcome switches (correct→error or error→correct) and those following outcome repetitions (correct→correct or error→error). The results for CNs and ENs were shown in red and blue, respectively. Data from two monkeys were shown in **a** and **b** separately. The black dot denotes the time point where there was significant difference between the outcome switch and outcome repetition trials (*: $P < 0.01$, paired t-test).

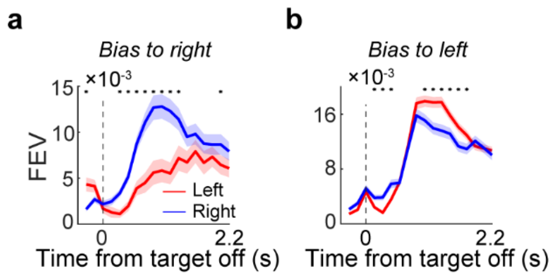


**Supplementary Fig. 14 Outcome selectivity in 7a correlated with monkeys' choice bias during AL. a-b**. Comparisons of outcome selectivity between the left saccade and right saccade trials in the ISAL task. Data sessions in which monkeys exhibited different saccade direction bias were shown separately. The outcome selectivity for different saccade directions were quantified separately for each neuron. Neurons recorded from different recording sessions were pulled together based on monkeys' saccade choice bias in the corresponding recording session. Stars denote the time points with significant difference between CNs and ENs ($P < 0.01$, paired t-test). The error bar denotes SEM.

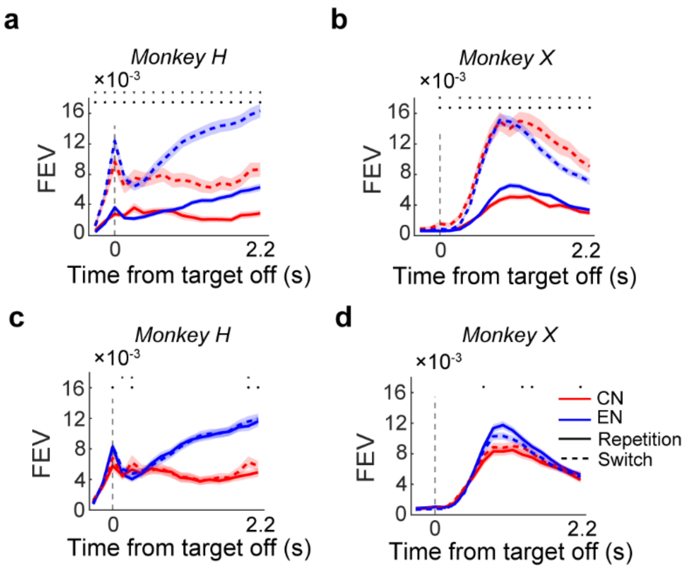


**Supplementary Fig. 15 Outcome representations in area 7a directly contribute to exploratory choice during learning, rather than reflect general motor variability. a-b**. Comparison of the averaged outcome selectivity between trials preceding exploratory choices (monkeys switched saccade direction relative to the previous trial for the same image–saccade association) and trials preceding exploitative choices (monkeys repeated the same saccade direction for the same association). Data from two monkeys are shown in the left and right panels separately. **c-d.** Comparison of the averaged outcome selectivity between trials preceding saccade switches and saccade repetitions, regardless of the image–saccade association. The black dot denotes the time point where the outcome encoding was significantly stronger in trials prior to switching choice than repetition choice ($P < 0.01$, paired t-test).

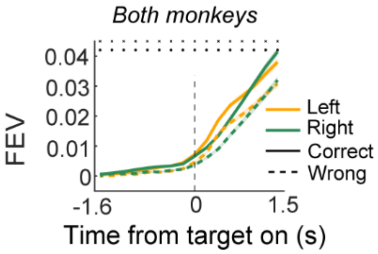


**Supplementary Fig. 16 Outcome history modulated the encoding of image–saccade associations.** Shown is a comparison of saccade direction encoding (quantified by FEV) between trials preceded by correct versus incorrect outcomes. Data for left-preferring and right-preferring neurons are presented separately. Solid and dashed lines represent trials following incorrect and correct outcomes, respectively. Black dots indicate time points with significant differences between the two conditions (paired t-test, $P < 0.01$). Shaded error bars represent the SEM.

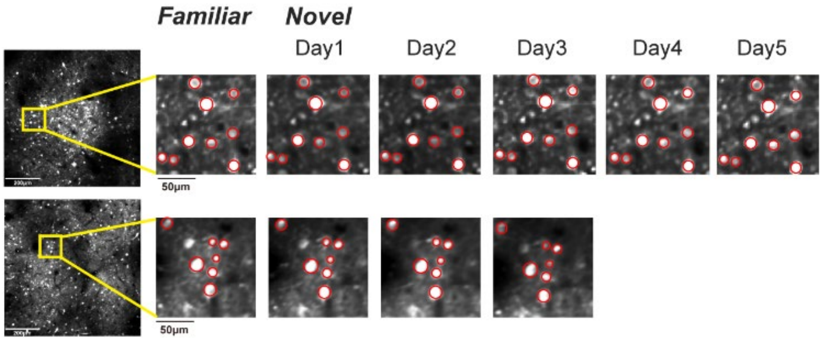


**Supplementary Fig. 17 Consistent identification of neurons across different learning days.** The left panels show the average two-photon images from two example fields of view (FOVs). The right panels display magnified sections of these FOVs, highlighting different recording days from two representative learning sessions. Yellow rectangles denote the regions magnified within each FOV, while red circles indicate neurons that were consistently identified within these regions across multiple days of the learning process.

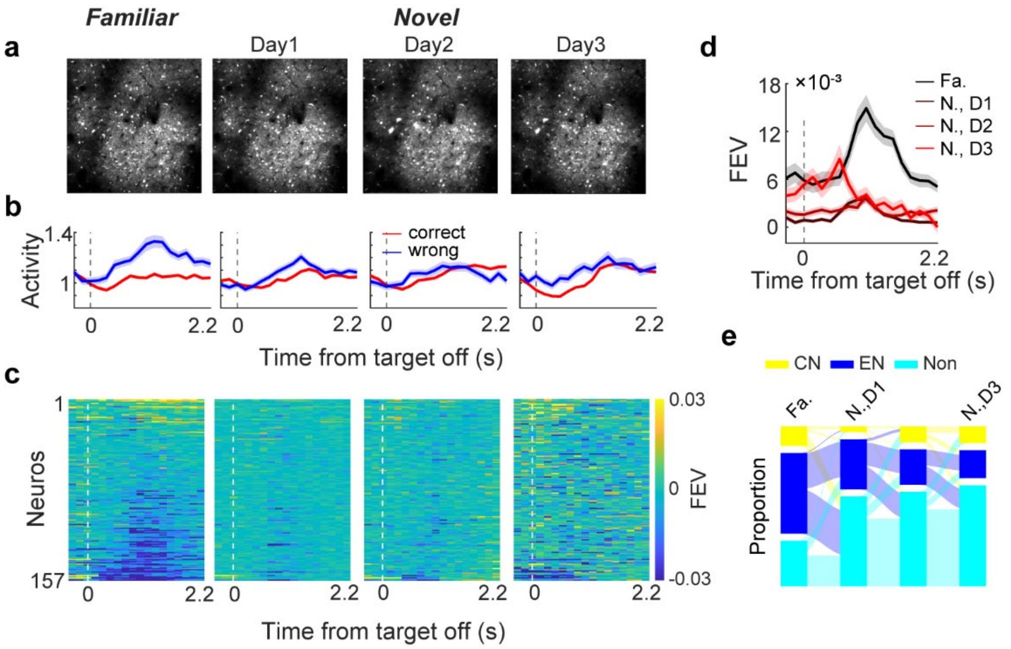


**Supplementary Fig. 18 Evolution of outcome selectivity in 7a during AL. a.** The average two-photon images of an FOV for different days within one example learning course. we tracked the activities of 157 neurons across four different sessions\days. **b.** An example neuron in this FOV. This neuron exhibited significant outcome encoding during the familiar day; but its outcome selectivity vanished after monkey transitioned to learn novel associations **c.** Outcome selectivity of all neurons identified within this example FOV were shown for different days within the learning course. Neurons in different days were listed in the same order. **d.** The averaged outcome selectivity of all neurons identified in this example FOV. **e.** Changes in cell types in this FOV across different days within the learning course.

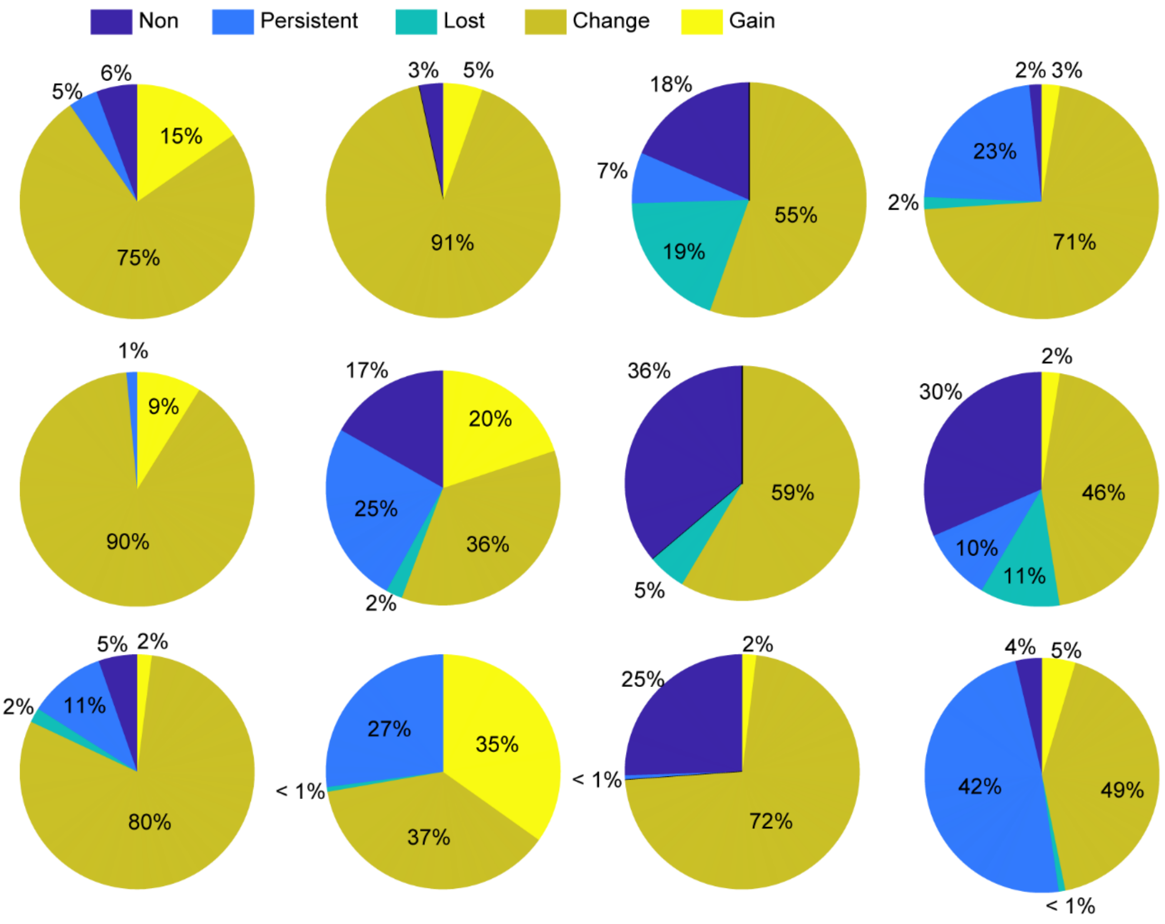


**Supplementary Fig. 19 The outcome encoding dramatically changed during the sensorimotor AL across all tested FOVs.** Each pie plot shows the result from one learning course (FOV). Different colors in the pie plots denote the proportions of neurons that changed outcome selectivity in different ways during AL. Non: non-selective; Persistent: persistently showed similar selectivity; Lost: lost selectivity; Change: changing selectivity; Gain: gain selectivity.

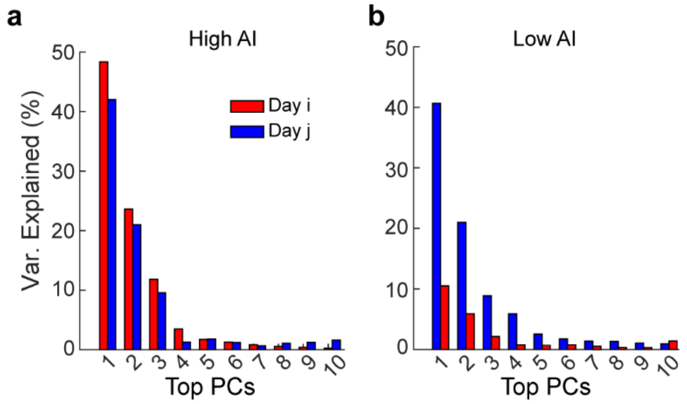


**Supplementary Fig. 20 Illustration of alignment index (AI) calculation.** This figure illustrates the process of calculating the Alignment Index (AI), a metric used to quantify the degree of alignment or overlap between the neural subspaces of population activity from different days. AI provides insights into how neural subspaces encoding trial outcomes evolve across learning days. To calculate the AI between day i and day j, we performed principal component analysis (PCA) on activity matrix from day i to obtain the PCs of day i. We selected the top 10 PCs to form a 10-dimensonal orthogonal axis in N-dimensional neural space. The overlap between neural subspaces (i.e. AI) can be estimated by projecting the activity matrix from day j onto the PCs of day i. It quantifies the proportion of variance explained relative to the total variance of population activity in day j. **a.** An illustration of the percentage of variance of data from day i explained when projected onto its own top10 PCs (red) or onto the top10 PCs defined by activity from day j (blue). In this case, the population codes between two days are highly overlapped. Thus, the resulting AI value is high. **b.** An example of low AI value.

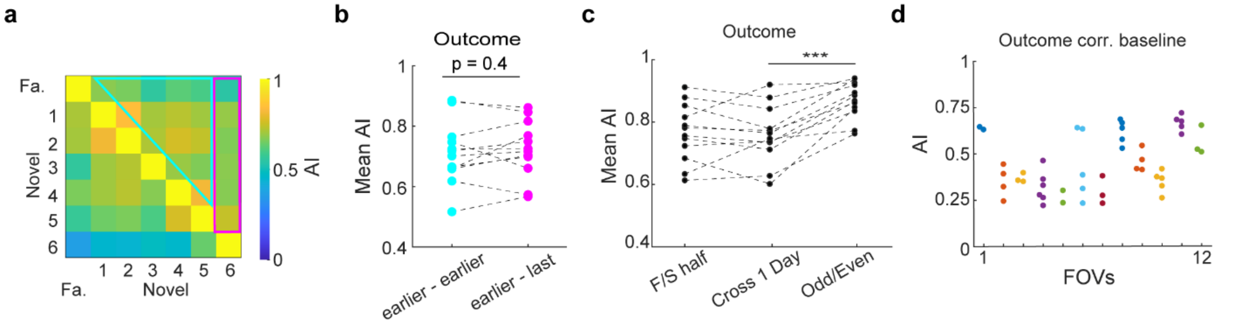


**Supplementary Fig. 21 Examining the influence of different factors on outcome encoding changes during AL.** We used alignment index to quantified the similarity of population codes for outcome encoding between different days. **a-b.** A control analysis showing that the remapping of the population-level outcome encoding after monkeys transitioned to learn new associations resulted from neither the difference in monkeys' behavior performance nor the difference in the temporal gaps between familiar day and learning days. In Figure 4M and 4O, we divided the data into novel-familiar and novel-novel groups, revealing that the alignment indices (AIs) for outcome encoding in the novel-familiar group were significantly smaller than those in the novel-novel groups. This indicated a reorganization of population-level outcome encoding after monkeys transitioned to learn new associations. However, a potential confounding factor in that analysis might be that the differences in monkeys' behavioral performance and temporal gaps were not balanced between the two groups. To address this, we further divided the AIs into two additional groups, ensuring they had similar levels of differences in behavioral performance and temporal gaps as those in the novel-familiar and novel-novel groups. **a.** We defined the familiar day and learning day in reverse sequence compared to Figure 4M. The last day in each learning course was defined as the familiar day, while the remaining days in each learning course were defined as learning days. Consequently, we divided the AIs into earlier-earlier and last-earlier groups. **b.** A comparison of the AIs between the earlier-earlier and last-earlier groups is presented, with each data point representing the averaged result for one FOV. Dashed lines connect data points from the same learning course. **c-d**. We investigate whether the evolution of outcome encoding during AL primarily arise from variations in the signal-to-noise ratio across different days. **c.** The AI values for outcome encoding at different time scales are depicted. To assess whether the changes in outcome encoding during AL were a result of variations in the signal-to-noise ratio across days, we calculated AIs to quantify the stability and similarity of population-level outcome encoding within single days and across different days. The neural data from each day were divided into two equal-sized parts based on the trial sequence (e.g., first and second halves of all trials or odd and even trials), and AIs were computed using these different subsets of data. (***: $P < 0.001$, paired t-test). **d.** The AIs here quantify the correlation between changes in outcome selectivity and changes in baseline activity (during the fixation period). Each data point represents data from two consecutive days, with different colors indicating different learning courses. The majority of AIs hover around 0.5, suggesting that there is no consistent correlation between changes in outcome encoding and baseline activity in 7a neurons during AL.

| | Region | | | | | | | | | | | |
|---|---|---|---|---|---|---|---|---|---|---|---|---|
| **Recording day** | **1** | **2** | **3** | **4** | **5** | **6** | **7** | **8** | **9** | **10** | **11** | **12** |
| **1** | 124 | 119 | 157 | 157 | 151 | 131 | 152 | 200 | 150 | 155 | 195 | 152 |
| **2** | 124 | 119 | 156 | 157 | 151 | 131 | 152 | 200 | 150 | 155 | 195 | 148 |
| **3** | 124 | 119 | 157 | 157 | 150 | 131 | 152 | 200 | 150 | 155 | 174 | 143 |
| **4** | | 119 | 149 | | 149 | | 152 | 200 | 150 | 155 | 182 | 133 |
| **5** | | | | | 150 | | 152 | | 150 | 155 | 171 | 126 |
| **6** | | | | | 138 | | 152 | | 150 | | 175 | 128 |
| **7** | | | | | 150 | | | | | | | 112 |
| **Cross day** | 124 | 119 | 148 | 157 | 135 | 131 | 152 | 200 | 150 | 155 | 157 | 109 |
| **Proportion (%)** | 100 | 100 | 95.6 | 100 | 91 | 100 | 100 | 100 | 100 | 100 | 86.2 | 81 |

**Supplementary Table 1 Number of neurons consistently identified across different recording days within each FOV.**

| Association pair | Learning day | |
|---|---|---|
| | Monkey H | Monkey X |
| 1 | 61 | 49 |
| 2 | 3 | 7 |
| 3 | 3 | 3 |
| 4 | 3 | 2 |
| 5 | | 6 |
| 6 | | 2 |
| 7 | | 5 |
| 8 | | 3 |
| 9 | | 5 |
| 10 | | 4 |
| 11 | | 5 |
| 12 | | 6 |

**Supplementary Table 2 Number of days monkeys spent learning each pair of novel associations.**
